# Systems-wide dissection of organic acid assimilation in *Pseudomonas aeruginosa* reveals a novel path to underground metabolism

**DOI:** 10.1101/2022.08.25.505149

**Authors:** Stephen K. Dolan, Andre Wijaya, Michael Kohlstedt, Lars Gläser, Paul Brear, Rafael Silva-Rocha, Christoph Wittmann, Martin Welch

## Abstract

The human pathogen *Pseudomonas aeruginosa* (Pa) is one of the most frequent and severe causes of nosocomial infection. This organism is also a major cause of airway infections in people with cystic fibrosis (CF). Pa is known to have a remarkable metabolic plasticity, allowing it to thrive in diverse environmental conditions and ecological niches, yet little is known about the central metabolic pathways which sustain its growth during infection, or precisely how these pathways operate. In this work, we used a combination of ‘omics approaches (transcriptomics, proteomics, metabolomics and ^13^C-fluxomics) and reverse genetics to provide a systems-level insight into how the infection-relevant organic acids, succinate and propionate, are metabolized by Pa. Moreover, through structural and kinetic analysis of the 2-methylcitrate synthase (PrpC) and its paralogue, citrate synthase (GltA), we show how these two crucial enzymatic steps are interconnected in Pa organic acid assimilation. We found that Pa can rapidly adapt to the loss of GltA function by acquiring mutations in a transcriptional repressor, which then de-represses *prpC* expression. Our findings provide a clear example of how ‘underground metabolism’, facilitated by enzyme substrate promiscuity, “rewires” Pa metabolism, allowing it to overcome the loss of a crucial enzyme. This pathogen-specific knowledge is critical for the advancement of a model-driven framework to target bacterial central metabolism.

## Introduction

*Pseudomonas aeruginosa* (Pa) is a notorious opportunistic human pathogen that frequently infects the airways of people with cystic fibrosis (pwCF). Pa is also well known for being metabolically flexible. This flexibility is important because nutrient acquisition and assimilation during infection scenarios is likely to be complex and dynamic process. Indeed, there is an increasing realisation that the enzymes of metabolism may also serve as targets for the next generation of antimicrobial therapies (1). However, we currently lack a clear understanding of how core metabolism operates in Pa.

Laboratory strains of Pa are known to prefer C4-dicarboxylates such as malate, fumarate and succinate as carbon and energy sources during growth *in vitro* (2). However, for reasons that are not yet clear, during infection, Pa frequently uses less-favoured carbon sources for growth, including the host-derived airway surfactant, phosphatidylcholine (PC). This phospholipid is broken down by secreted Pa phospholipases to yield phosphorylcholine, glycerol, and long-chain fatty acids (3, 4).

Pa can also metabolize short chain fatty acids. Propionate is a naturally-occurring short-chain fatty acid produced by the human gut microbiota, and is a commonly used food preservative with potent bacteriostatic activity. Another rich source of propionate are the anaerobes that frequently occupy in the lower airways of pwCF. These anaerobes break down tracheobronchial mucin to produce copious quantities of propionate. However, and in spite of its known growth-inhibitory properties against some species of bacteria, Pa is able to thrive on propionate, and can very effectively utilise the compound as a sole carbon source *in vitro* (5–7). Pa does this by catabolising propionate through the 2-methylcitrate cycle (2MCC) (Figure 1A) to yield succinate and pyruvate, which feed directly into the TCA cycle. The 2MCC also sits at an important junction in amino acid catabolism, since several amino acids (L-valine, L-*iso*leucine, L-methionine, and L-threonine) are degraded to propionyl-CoA, which must then be oxidised by this pathway (7). Given the ubiquity of propionate in many host niches, it comes as little surprise that a functional 2MCC is required for infection by a plethora of human pathogens, including *Mycobacterium tuberculosis*, *Neisseria meningitides* and *Aspergillus fumigatus* and *Talaromyces marneffei* (8–12). The Pa 2MCC has also been shown to be important for infection of the nematode intestine (13).

**Figure 1.**
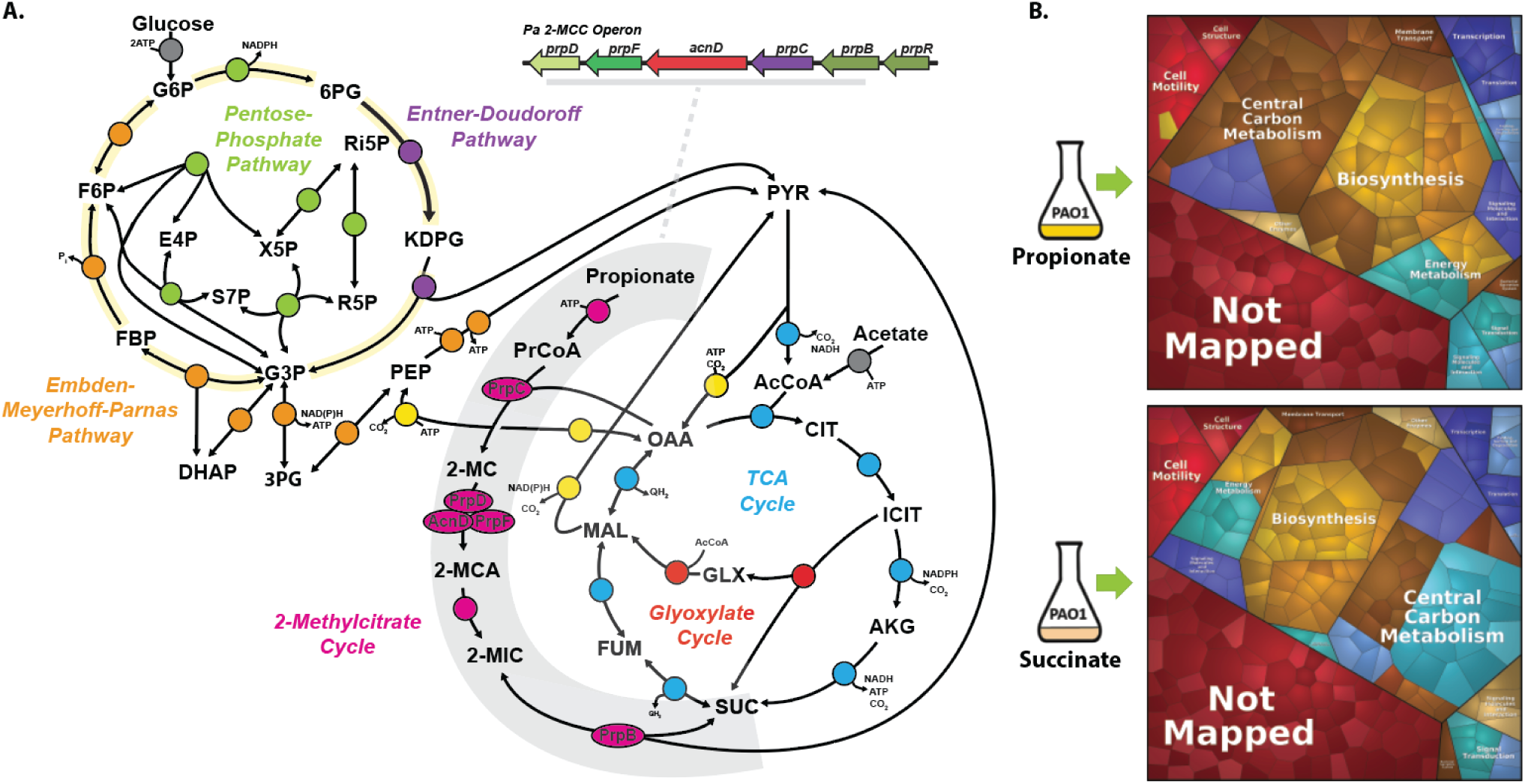
Proteomic analysis of Pa grown on Succinate and Propionate. A; Schematic depicting the Pa 2-methylcitrate cycle (2MCC) in Pa central carbon metabolism. The Pa central metabolic network shown here consists of six main blocks, designated with different colours: (i) the Embden-Meyerhoff-Parnas pathway (EMP, orange); (ii) the pentose phosphate pathway (PPP, green); (iii) the Entner-Doudoroff pathway (EDP, purple); (iv) the tricarboxylic acid cycle (TCA, blue) and glyoxylate shunt (red); (v) anaplerotic and gluconeogenic reactions (yellow), and (vi) the 2MCC (pink). The 2MCC operon arrangement (inset – grey underline) consists of genes which encode; a transcriptional regulator (designated here as *prpR*) which is thought to encode a ligand- responsive repressor, a methylcitrate synthase (*prpC*) which condenses propionyl-CoA (PrCoA) with oxaloacetate (OAA) to form 2-methylcitrate (2-MC), a 2-methylcitrate dehydratase/hydratase (*prpD*) which dehydrates 2-MC to yield 2-methylaconitate (2-MCA), a 2-methylcitrate dehydratase (*acnD*) and 2-methylaconitate *cis-trans* isomerase (*prpF*) which provide an alternative route for the generation of 2-MCA from 2-MC, and a 2- methyl*iso*citrate lyase (*prpB*), which cleaves 2-methyl*iso*citrate (2-MIC) to yield pyruvate (PYR) and succinate (SUC). Note that the 2-MCA generated in the PrpD or AcnD/PrpF reactions is rehydrated by an unlinked aconitase (likely AcnB in Pa) to yield the PrpB substrate, 2-MIC. Also, the enzyme responsible for the initial activation of propionate to yield PrCoA has not yet been identified for Pa, although in other organisms this function is carried out by a dedicated propionyl-CoA synthase (PrpE), or by acetyl-CoA synthase (AcsA), or by a combination of phosphotransacetylase (Pta) and acetate kinase (AckA) activities. Other abbreviations; AcCoA, acetyl-coenzyme A; CIT, citrate; ICIT, *iso*citrate; AKG, α- ketoglutarate; FUM, fumarate; MAL, malate; KDPG, 2-keto-3-deoxy-6-phosphogluconate; G3P, glyceraldehyde 3-phosphate; FBP, fructose 1,6-*bis*phosphate; F6P, fructose 6- phosphate; G6P, glucose 6-phosphate; 6PG, 6-phosphogluconate; Ri5P, ribulose 5-phosphate; R5P, ribose 5-phosphate; X5P, xylulose 5-phosphate; S7P, sedoheptulose 7- phosphate; E4P, erythrose 4-phosphate; PEP, phospho*enol*pyruvate. B; Illustration of the statistically significant proteomic changes (p-value ≤ 0.05, fold change ≥1 or ≤-1) during growth on propionate or succinate, as represented by Voronoi tessellations. Pathway assignment was performed using the Kyoto Encyclopedia of Genes and Genomes (KEGG) dataset. Proteome alterations which could not be assigned to a specific pathway (uncharacterised/hypothetical proteins) are shown as ‘Not Mapped’. The specific protein identities for the protein clusters that were up-regulated during growth on propionate are shown in Figure S1A and statistical analyses of these data are illustrated in Figures S1B-D. The complete proteomics dataset is presented in File S1.

We currently have a limited understanding of how Pa metabolises propionate, or how the 2MCC interfaces with the other components of central carbon metabolism in this organism. Although some features of the pathway can be extrapolated from a knowledge of the biochemistry in other bacteria (such as *Escherichia coli* and *Salmonella enterica*) these are fundamentally dissimilar microbes with alternative operonic arrangements for the 2MCC ORFs, and very different metabolic architectures compared with Pa (14, 15). For example, propionate metabolism in several *Enterobacteriales* (including *E. coli* and *S. enterica*) and all analysed *Xanthomonadales* is coordinated by a Fis family transcription factor (TF) known as PrpR (16). By contrast, the 2MCC in *Gammaproteobacteria* is typically controlled by a GntR family TF. Remarkably, no 2MCC regulators from the GntR family have been experimentally characterised to date. Therefore, and to understand better how Pa utilizes propionate, we used a combination of ‘omics approaches (transcriptomics, proteomics, metabolomics and ^13^C-fluxomics) and reverse genetics to provide a systems-level insight into how the organic acids, succinate and propionate, are metabolized by Pa. Moreover, through structural and kinetic analysis of the 2-methylcitrate synthase (PrpC), and its paralogue citrate synthase (GltA), we show how these two crucial steps are interconnected in organic acid assimilation. Building on these observations, we found that Pa can rapidly adapt to the loss of GltA by acquiring mutations that de-repress expression of the *prpC*-encoding 2MCC operon (*prp*). These mutations are in a GntR-family TF, which we show encodes a transcriptional repressor of the *prp* operon. Our findings provide a clear example of how ‘underground metabolism’ (17), facilitated by enzyme promiscuity, allows Pa to overcome the loss of a crucial enzyme in central carbon metabolism.

## Results

### ‘Omics-driven examination of Pa grown on succinate and propionate as sole carbon sources

To understand how growth on different substrates affects the physiology of Pa, we first examined the proteome during exponential phase growth on succinate or on propionate as a sole carbon source. Through the proteomic analysis, we identified and quantified 3796 proteins. Of these, 265 proteins showed increased abundance during growth on propionate, and 295 proteins showed increased abundance during growth on succinate (*q*-value ≤ 0.05, log_2_ fold-change ≥1 or ≤-1; File S1). To obtain a global overview of the physiological changes, we used the Proteomaps web service (18) to generate Voronoi tessellations (19) structured around the KEGG orthologies of the statistically significant changes (*p*-value ≤ 0.01, log_2_ fold change ≥1 or ≤-1). As shown in Figure 1B, most of the proteomic changes were associated with ‘central carbon metabolism’, ‘biosynthesis’, ‘signalling and cellular process’ and ‘energy metabolism’. Notably, growth on propionate led to a strong induction (ca. 16-fold change (FC)) of all proteins encoded by the *prp* operon, including the GntR-family 2MCC operon regulator, PA0797, which we designate here as PrpR (Figure S1A-D, File 1A).

To provide a complementary insight into the absolute metabolic fluxes in Pa during growth on propionate and succinate, we also carried out a [^13^C] fluxome analysis. This was achieved by measuring the mass isotopomer distributions in proteinogenic amino acids and cell carbohydrates using three separate tracers for propionate and succinate (*Materials and Methods*) (20). The calculated relative fluxes for Pa strain PAO1 grown on labelled propionate or succinate are shown in Figure 2. The corresponding quantitative comparison of NADPH (redox) supply and ATP (energy) supply for succinate- and propionate-grown Pa are shown in Figure S2.

**Figure 2.**
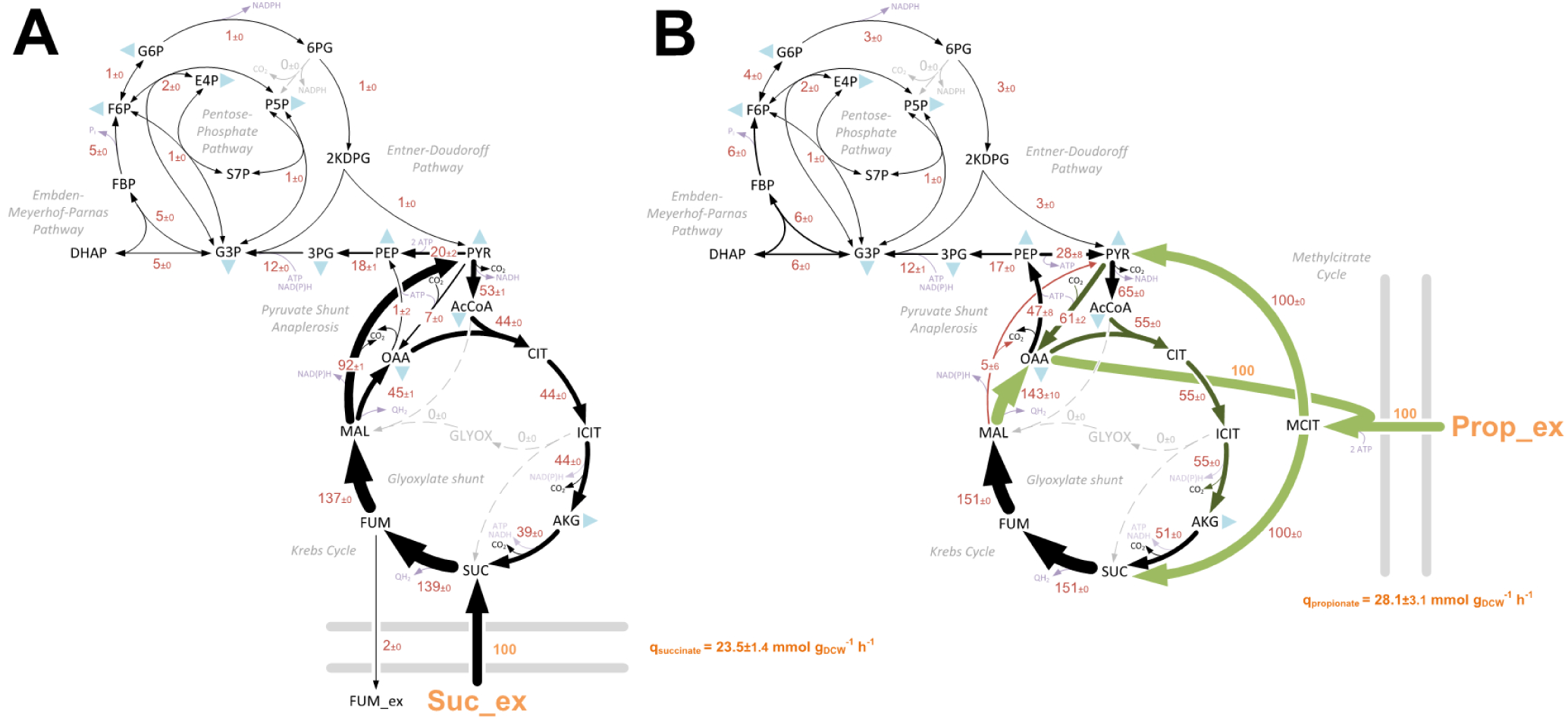
*In vivo* carbon flux distributions in central metabolism of Pa PAO1 during growth on succinate. (A) or propionate (B) as sole carbon sources. Flux is expressed as a molar percentage of the average uptake rate for succinate (23.5 mmol g^−1^ h^−1^) or propionate 28.1 mmol g^−1^ h^−1^), calculated from the individual rates in File S2. Anabolic pathways from 11 precursors to biomass are indicated by the filled blue triangles. The flux distributions with bidirectional resolution (i.e., net and exchange fluxes), including the drain from metabolic intermediates to biomass and confidence intervals of the flux estimates, are provided in File S2. The errors given for each flux reflect the corresponding 90% confidence intervals. The full flux datasets are presented in *Supporting Information* File S2. Colours qualitatively indicate fluxomic correlation with changes on the protein level during growth on propionate compared with growth on succinate (light green or red → significant up- or down-regulation (respectively); dark green or red → less significant up- or down-regulation).

Comparison of the flux maps, in combination with the proteomic data, generated an unparalleled insight into the central carbon metabolic networks of Pa during growth on both substrates. For example, several of the key proteomic alterations found when comparing growth on propionate with growth on succinate were in core central carbon metabolism (File S1A). These core changes were largely consistent with the corresponding carbon flux distributions (Figure 2). In general, the expression of enzymes from the pentose phosphate pathway (PPP), the Embden-Meyerhof-Parnas pathway (EMPP) and the Entner-Doudoroff pathway (EDP) was decreased during growth in propionate compared with succinate. The expression of several enzymes in the TCA cycle were increased during growth on propionate, including citrate synthase GltA (2.5 FC), aconitase AcnA (2.3 FC) and the *iso*citrate dehydrogenases ICD (2.5 FC) and IDH (1.7 FC). A corresponding increase in TCA cycle carbon flux was also evident, with roughly an 11% increase in flux through the reactions between citrate (CIT) and malate (MAL) (Figure 2). Fumarate efflux (2%) was also detected during Pa growth using succinate as a sole carbon source.

Among the largest discrepancies between the propionate- and succinate-grown cultures at both the proteome and fluxome level were noted at the reactions involved in the pyruvate shunt. Expression of the malic enzyme, MaeB, which catalyses the oxidative decarboxylation of malate to produce pyruvate and CO_2_, was down-regulated (-4.7 FC) during growth on propionate. By contrast, the pyruvate carboxylase-encoding proteins, *pycA* (PA5435) and *pycB* (PA5436), which catalyse the ATP-dependent carboxylation of pyruvate to yield oxaloacetate, were up-regulated (2.6 FC), as was the regulator, PycR (21, 22). These alterations matched the corresponding flux data, which revealed a substantial decrease in flux from malate to pyruvate (-87%) and an increase in the flux from pyruvate to oxaloacetate (OAA) (54%) during growth on propionate. There is a good metabolic logic to this. Although the catabolism of propionate yields succinate and pyruvate, an early enzyme in the propionate catabolic pathway (PrpC) requires oxaloacetate as a substrate (Figure 1). Therefore, we hypothesised that this drain on the oxaloacetate pool may be countered by a combination of lower malic enzyme-mediated pyruvate generation and increased anaplerotic pyruvate carboxylase activity to sustain the TCA cycle. In support of this, mutants defective in pyruvate carboxylase (encoded by *pycA* and *pycB*) were unable to grow on propionate as a sole carbon source (Figure S1E). Growth on propionate also increased the expression (5.1 FC) of the membrane-bound malate-quinone oxidoreductase, MqoB, which generates oxaloacetate directly from malate. A corresponding 98% increase in carbon flux from malate to oxaloacetate was evident during growth on propionate (Figure 2).

The expression level of phospho*enol*pyruvate synthase (PPS) and pyruvate kinase showed no significant differences between the growth conditions. However, the fluxomic analysis captured a pronounced alteration in carbon flow at this node. During growth on succinate, the net carbon flux was in the pyruvate → phospho*eno*lpyruvate (gluconeogenic) direction, with phospho*eno*lpyruvate originating from pyruvate mainly *via* the combined activity of malic enzyme (MAE) and PEP synthase at the equivalent cost of 2 ATP (PYR + H_2_O + ATP → PEP + AMP + P_i_). By contrast, during growth on propionate, the net flux at this node was in the direction phospho*eno*lpyruvate → pyruvate; a reaction that is catalyzed by pyruvate kinase isozyme A (PykA) and that generates ATP (23). In this scenario, phospho*eno*lpyruvate largely originates from the action of phospho*enol*pyruvate carboxykinase (PckA) on the oxaloacetate that is generated *via* the activity of malate-quinone oxidoreductase (MqoB). Interestingly, and despite the greatly increased flux from oxaloacetate → phospho*enol*pyruvate catalyzed by PckA, the expression of this enzyme was decreased 1.7-fold during growth on propionate compared with growth on succinate. This may indicate a role for allosteric regulation in modulating PckA enzyme activity (24, 25).

When compared with growth on succinate, the glyoxylate shunt enzymes, *iso*citrate lyase (AceA; 5.5 FC) and malate synthase (GlcB; 3.6 FC) were highly expressed on propionate. This was also verified using promoter-luciferase transcriptional fusions (Figure S1F-I). However, the fluxomics data indicated that there was <u>no</u> carbon flux through the glyoxylate shunt during growth on either succinate or propionate as a sole carbon source. This may be explained by the extensive allosteric interactions which are known to control flux partitioning between the TCA cycle and glyoxylate shunt. *Iso*citrate lyase (ICL) activity in Pa is allosterically inhibited by oxaloacetate, pyruvate, succinate, phospho*enol*pyruvate (PEP), and CoA. By contrast, oxaloacetate and pyruvate allosterically activate one of the *iso*citrate dehydrogenase enzymes, IDH (26). As flux to pyruvate is significantly increased during growth on either succinate or propionate compared with acetate (where flux through the glyoxylate shunt is maximal), these data suggest that pyruvate is the most likely metabolite responsible for abrogating flux through the glyoxylate shunt during growth on propionate (27).

Several studies have suggested that simple diffusion across the cytoplasmic membrane is the predominant mechanism of both acetate and propionate uptake in bacteria (28, 29). One cluster of ORFs (PA3232-PA3235) were up-regulated (ca. 16 FC) during growth on propionate, and encode a putative acetate permease (ActP). Indeed, PA3234 shows 80% amino acid identity to the ActP protein from *Escherichia coli*, and this ORF has been previously shown to be regulated by the two component system, MxtR*/*ErdR, which is essential for growth on acetate (30, 31). MxtR was also up-regulated during growth on propionate (8.7 FC), whereas proteins associated with dicarboxylic acid transport (DctA, DctQ, DctP and PA5530) were more abundant during growth on succinate (File S1A) (32, 33).

Aerobic growth in different carbon sources results in large-scale remodelling of the electron transport chain in Pa, including components of the denitrification pathway (27). Growth on propionate led to significantly increased expression of most terminal oxidases, particularly the quinol oxidase Cyo (7.6 FC), the cyanide-insensitive oxidase Cio (5.4 FC), the cytochrome c oxidase Cco2 (3.0 FC) and the cytochrome oxidase Cox (2.2 FC). Furthering the notion of an altered redox balance during growth on the two substrates, we noticed differences in expression of the NAD(P) transhydrogenases, which fine-tune the size and degree of reduction of the nicotinamide adenine dinucleotide pools (34). Expression of the transhydrogenase Sth (PA2991) which is thought to primarily convert NADPH → NADH, was increased during growth on propionate (2.1 FC), whereas the transhydrogenase proteins PntAA (3.6 FC) and PntB (2.2 FC) (which presumably catalyze the interconversion NADH → NADPH) were more abundant during growth on succinate. These alterations were reflected in the redox balances; growth on propionate resulted in a lower NADPH surplus (as any excess is assumed to be converted to NADH to drive ATP synthesis) when compared with growth on succinate (Figure S2).

The 2MCC also serves a role in the catabolism of branched chain amino acids (BCAA). This is because *iso*leucine and valine degradation generates propionyl-CoA, which can then be degraded to succinate and pyruvate *via* the 2MCC (35). The metabolism of valine produces the intermediate (S)-3-hydroxy*iso*butyric acid which is oxidized to methylmalonate semialdehyde by 3-hydroxy*iso*butyrate dehydrogenase (MmsB). Methylmalonate semialdehyde dehydrogenase (MmsA) then catalyses the irreversible NAD^+^- and CoA-dependent oxidative decarboxylation of the semialdehyde to yield propionyl-CoA (35, 36). MmsB (28.1 FC) and MmsA (5.3 FC) expression were significantly increased during growth on propionate. This suggests that there is a regulatory link between the 2MCC and BCAA catabolism in Pa.

### Propionate inhibits the growth of Pa when propionate catabolism is disrupted

Based on the proteomics data, we made (separate) in-frame deletions in a selection of genes putatively involved in propionate uptake (PA3234 – the *actP* homologue), propionate activation (*acsA*, PA3568) and propionate catabolism (*prpC*, *mmsA*, *aceA*, *glcB*) and tested the ability of the resulting mutants to grow on a series of single carbon sources (Figure 3A, Figure S3A-C). Importantly, the acetyl-CoA synthetase mutant (Δ*acsA*) exhibited a pronounced growth defect with either acetate or propionate as a sole carbon source, suggesting that activation of propionate is carried out primarily by this enzyme (Figure S3B,C). By contrast, the ΔPA3568 mutant (defective in another potential propionyl-CoA synthetase) and the putative acetate symporter (ΔPA3234) mutant had no growth phenotype in any of the conditions tested. Similarly, the mutants defective in *mmsA*, *aceA*, or *glcB* also exhibited no growth defects on propionate, despite significant up-regulation of the corresponding gene products during growth on this carbon source. Notably, the 2- methylcitrate synthase mutant (Δ*prpC*) was unable to grow on propionate as a sole carbon source but displayed no detectable growth deficit on any of the other tested carbon sources.

**Figure 3.**
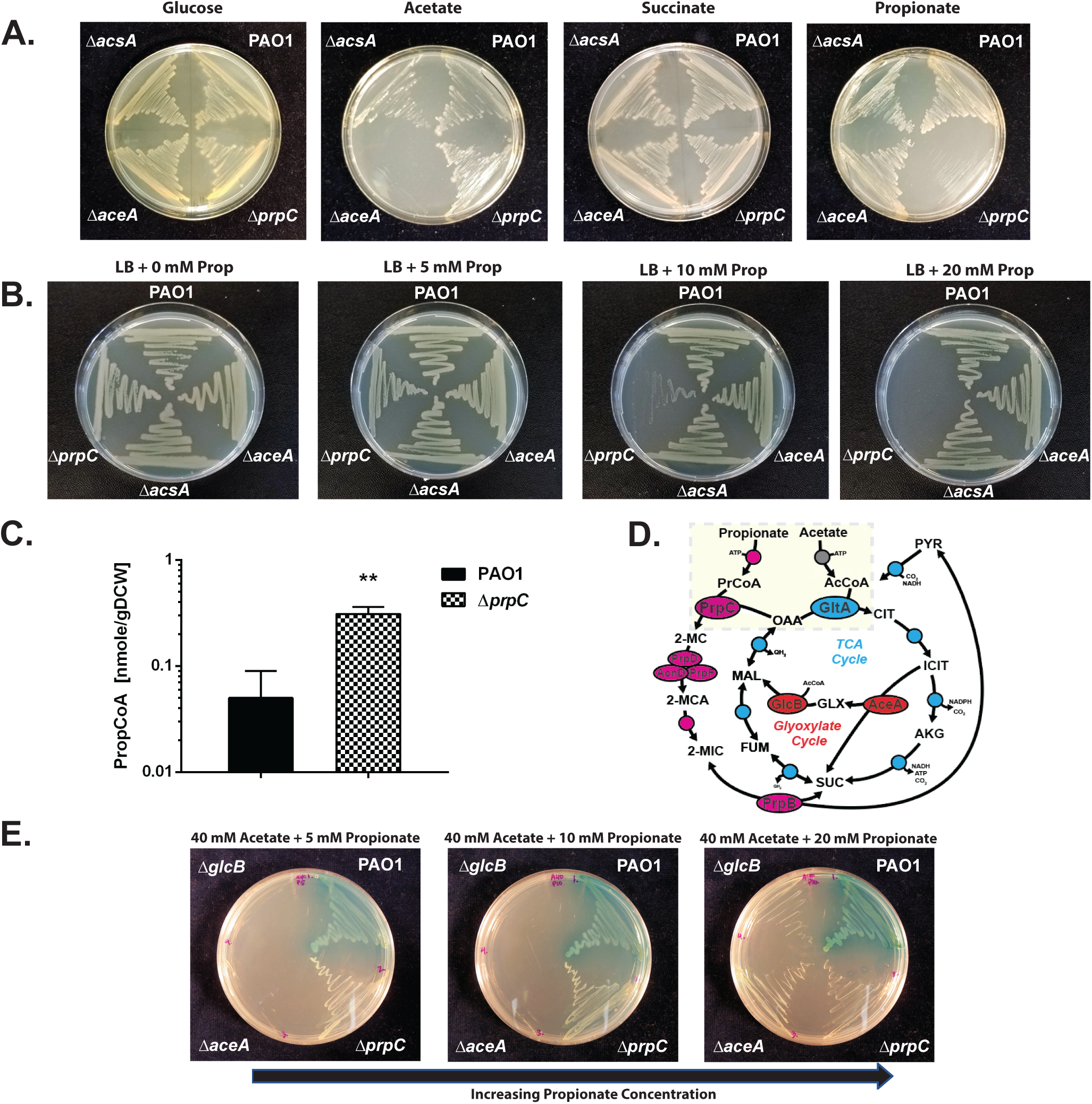
The Pa ORF (*prpC*) encoding 2-methylcitrate synthase is essential for growth on propionate. A; Wild-type Pa (PAO1), and the Δ*acsA*, Δ*aceA* and Δ*prpC* mutants all grow comparably on MOPS agar containing glucose or succinate as a sole carbon source. Δ*acsA* has a growth defect during growth on MOPS-acetate and MOPS-propionate. Δ*prpC* cannot grow on MOPS propionate. The plates were photographed after 24 h incubation. B; Wild- type PAO1 and the Δ*acsA*, Δ*aceA* and Δ*prpC* mutants were cultured on LB agar containing an increasing concentration of propionate (0, 5, 10, 20 mM, as indicated). The Δ*prpC* displays a pronounced growth defect in the presence of propionate concentrations > 10 mM. The plates were photographed after 24 h incubation. C; Intracellular propionyl-CoA concentration in wild-type Pa (PAO1) and in the Δ*prpC* mutant following 3h exposure to propionate (5 mM). Unpaired *t*-test with Welch’s correction p = 0.0026. The experiment was performed using biological triplicates. D; Illustration of the interwoven reactions for propionate and acetate activation in Pa, feeding into the 2-methylcitrate cycle and TCA cycle, respectively. Following uptake, acetate and propionate are activated by AcsA. The resulting propionyl-CoA (PrCoA) is condensed with oxaloacetate (OAA) in a PrpC-catalyzed reaction to form 2-methylcitrate (2-MC), whereas the acetyl-CoA (AcCoA) is condensed with oxaloacetate in a GltA-catalyzed reaction to form citrate (CIT). E; Growth of the Pa glyoxylate shunt mutants Δ*aceA* and Δ*glcB* is blocked on MOPS agar plates containing a combination of high acetate concentration (40 mM) and low propionate concentration (5 mM) as the carbon source. However, this growth inhibition is partially overcome by increasing the propionate concentration to 20 mM (left to right in the figure). The plates were photographed after 48 h growth. The data are representative of two independent experiments, each performed in triplicate.

In the absence of a functional 2-methylcitrate synthase (PrpC), the accumulation of intracellular propionyl-CoA, either through direct catabolism of propionate or through catabolism of branched chain amino acids, has been proposed to have growth inhibitory properties for several microorganisms (37–39). As shown in Figure 3B, when propionate (5- 20 mM) is added to LB agar, growth of the Δ*prpC* mutant becomes progressively more inhibited as the concentration of propionate increases. The growth inhibitory effect of propionate on the Δ*prpC* mutant was also apparent when propionate was added to MOPS-buffered succinate, ruling out pH dependent toxicity (Figure S3D). Notably, the Δ*prpC* mutant was also unable to grow on branched chain amino acids as a sole carbon source (Figure S3E). To examine this possible growth inhibition by propionyl-CoA further, we exposed PAO1 and the Δ*prpC* mutant to 5 mM propionate during exponential growth on succinate (25 mM). Then, after a further 3 h of growth, we measured the intracellular propionyl-CoA levels in each sample (Figure 3C). This revealed that even during growth on a preferred carbon source, succinate, propionyl-CoA accumulates in the Δ*prpC* mutant compared with wild-type PAO1.

We previously characterized the metabolic pathways expressed in Pa during growth on acetate (27). Comparison of those data with the results presented here for growth on propionate revealed several commonalities, including increased expression of AcsA, PA3234 (the *actP* homologue), and the glyoxylate shunt enzymes on acetate and propionate. This may reflect the activity of shared regulators, or analogous reaction mechanisms and overlapping substrates (Figure 3D). As shown in Figure S3F and consistent with the fluxomics data (which revealed negligible flux through the glyoxylate shunt during growth on propionate), mutants defective in the glyoxylate shunt enzymes, Δ*aceA* and Δ*glcB*, suffered no growth defect on propionate as a sole carbon source. As expected, the same mutants were unable to grow on acetate as a sole carbon source (Figure S3F). Remarkably, this growth of the Δ*aceA* and Δ*glcB* mutants on propionate was blocked when acetate was added to the media (Figure 3E). This growth inhibition could be partially relieved by increasing the concentration of propionate in the media, suggesting metabolic competition between acetate and propionate catabolism (Figure 3E). By contrast, acetate did not prevent growth of the Δ*aceA* and Δ*glcB* mutants on plates containing succinate (Figure S3F). In the absence of the glyoxylate shunt, acetyl-CoA generated through the activation of acetate or through β-oxidation of fatty acids is unable to contribute to Pa biomass generation. Since AcsA likely activates both acetate and propionate, a parsimonious hypothesis is that saturating concentrations of acetate (which is probably the preferred substrate of AcsA) competitively block the activation of propionate. This competition is relieved at higher propionate concentrations, thereby enabling growth of the Δ*aceA* and Δ*glcB* mutants.

### Structural and functional investigation of PrpC and GltA from Pa

PrpC catalyzes the condensation of oxaloacetate and propionyl-CoA. In a parallel reaction, the TCA cycle enzyme, citrate synthase (GltA), catalyzes the condensation of oxaloacetate and acetyl-CoA. Given the apparent promiscuity of AcsA with respect to acetate and propionate activation, we wondered whether the condensation of propionyl-CoA and acetyl-CoA with oxaloacetate could be carried out interchangeably by PrpC and GltA (Figure 3D). Indeed, PrpC from *E. coli* has secondary citrate synthase activity and overexpression of *prpC* in this organism can rescue the synthetic lethality of citrate synthase loss (40–43). To examine whether this is also the case in Pa, a Δ*gltA* mutant was generated. Colonies of the Δ*gltA* mutant on LB agar were visibly smaller when compared with wild-type PAO1 (Figure 4A). This phenotype could be partially complemented by supplementing the plates with glutamate, whose carbon skeleton enters the TCA cycle after the citrate synthase-catalyzed step (Figure S4A).

**Figure 4.**
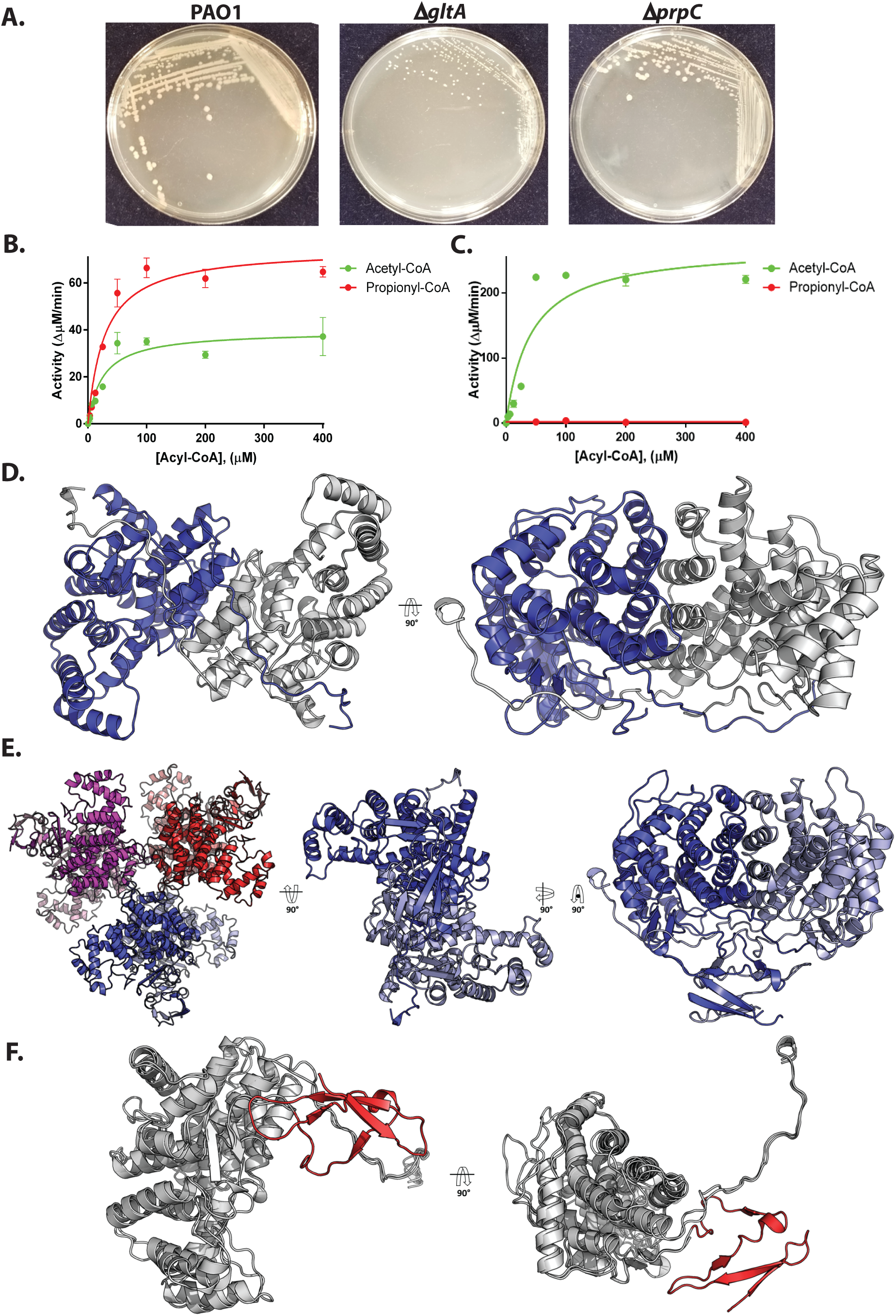
Biochemical and structural analysis of PrpC and GltA from Pa. A. A Δ*gltA* mutant exhibits a growth defect when cultured on LB-agar, whereas a Δ*prpC* mutant displays a wild-type colony morphotype. The plates were photographed after 48 h. The data are representative of two independent experiments, each performed in triplicate. B. Purified PrpC_Pa_ exhibits both citrate synthase activity (with acetyl-CoA as a substrate) and 2-methylcitrate synthase activity (with propionyl-CoA as a substrate). The concentration of OAA in each reaction was fixed at 0.5 mM. The data are representative of two independent experiments, each performed in triplicate. C. Purified GltA_Pa_ is a citrate synthase with no detectable 2-methylcitrate synthase activity. The concentration of OAA was fixed at 0.5 mM. The data are representative of two independent experiments, each performed in triplicate. D. The x-ray crystal structure of PrpC_Pa_ (PDB: 6S6F). PrpC_Pa_ is a homodimer. In the ribbon diagram shown, the protomers are coloured blue and grey. E. Cartoon representation of the GltA_Pa_ hexamer in the asymmetric unit (left) and, for comparison with PrpC_Pa_, the extracted GltA_Pa_ dimers (middle and right). F. Superposition of the PrpC_Pa_ and GltA_Pa_ structures. PrpC_Pa_ and GltA_Pa_ share similar core α- helical folds (shown in grey to highlight similarities). However, GltA_Pa_ has an additional antiparallel β-sheet at its N-terminus (coloured in red to showcase differences).

To assess directly whether PrpC_Pa_ has citrate synthase activity (and whether citrate synthase may also have 2-MC synthase activity), we purified each enzyme to investigate its specificity and kinetic properties *in vitro*. The Pa *prpC* and *gltA* genes were cloned and overexpressed (with cleavable His_6_ tags) in *E. coli*, and purified to homogeneity. Each purified enzyme was then assayed for 2-methylcitrate synthase activity and citrate synthase activity. The PrpC enzymes from species including *S. enterica*, *E. coli*, and *B. subtilis* have previously been reported to exhibit a strong preference for propionyl-CoA compared with acetyl-CoA (42, 44, 45). However, PrpC_Pa_ displayed roughly comparable activity towards these acyl-CoAs, although V_max_ was greater with propionyl-CoA as a substrate (Figure 4B, S4B). The specificity (expressed as k_cat_/K_M_) of PrpC_Pa_ for propionyl-CoA was 104^3^ x 10 M^-1^ s^-1^, whereas for acetyl-CoA k /K was 114 x 10^3^ M^-1^ s^-1^ (Table S1C). By contrast, and unlike GltA from *S. enterica* (which exhibits a low level of 2-methylcitrate synthase activity (46)), GltA from Pa (GltA_Pa_) had no detectable 2-methylcitrate synthase activity (Figure 4C).

To gain insights into the possible structural bases for these kinetic data, we used x- ray crystallography to solve the structure of PrpC_Pa_ and GltA_Pa_ (Figure 4D). PrpC_Pa_ is a homodimer in both the crystal structure (Figure 4D) and in solution (Figure S4D), whereas the GltA_Pa_ asymmetric unit was comprised of a hexameric “trimer of dimers” (Figure 4E). Structural superposition of PrpC_Pa_ and GltA_Pa_ revealed a near-identical α-helical core fold (Figure 4F) with a Cα RMSD of 1.33 Å. GltA_Pa_ is slightly larger than PrpC_Pa_ (429 amino acids *versus* 376 amino acids, respectively) and has an additional 50 amino acid residues at its N- terminus, which form four anti-parallel β-strands and loops (Figure 4F). In addition to solving the apo-structures of the enzymes, we also obtained the structure of PrpC_Pa_ with oxaloacetate bound in the active site (Figure 5A). The active site was located in a cleft between two domains on the enzyme. A comparison of PrpC structures from different bacterial species revealed that the residues comprising the PrpC_Pa_ active site are very highly conserved. For instance, in *S. enterica* His235 (His222 in PrpC_Pa_), His274 (His 261 in PrpC_Pa_), and Asp325 (Asp312 in PrpC_Pa_) form a catalytic triad (45).

**Figure 5.**
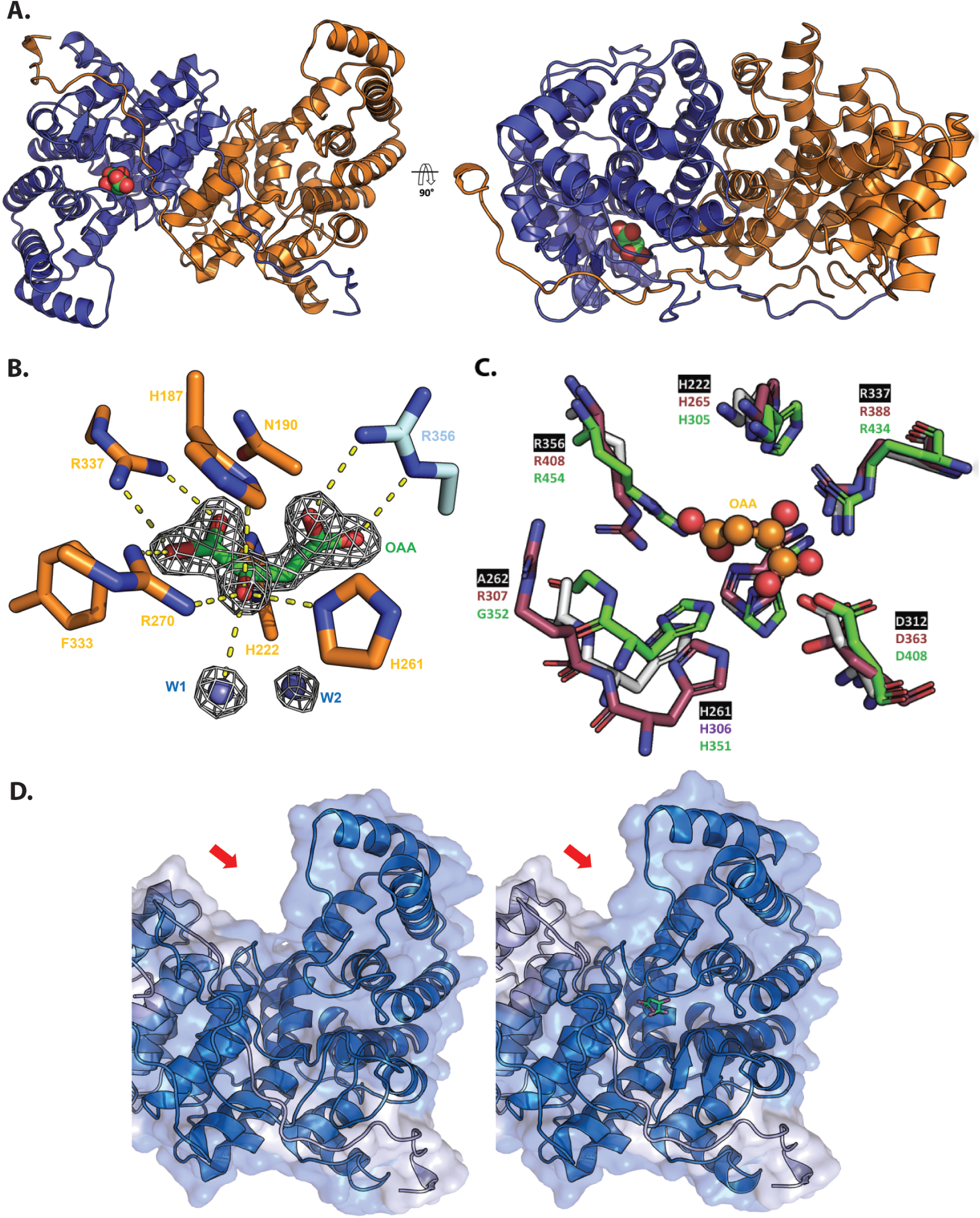
Structural analysis of oxaloacetate-bound PrpC_Pa_. A; Crystal structure of oxaloacetate-bound PrpC_Pa_ represented in cartoon (PDB: 6S87). One protomer is coloured blue and the other orange. A 90° rotation about the X-axis is shown (right). Oxaloacetate is shown as green and red spheres. B; Oxaloacetate binding site from *P. aeruginosa* PrpC chain D. Water molecules are shown in cyan spheres. Chain D and chain C residues are shown in orange and cyan, respectively. The electron density map (2*F*o*-F*c) in white is contoured at 1.5σ. C; Superposition of the PrpC_Pa_ (white), GltA_Pa_ (pink) and *A. fumigatus* PrpC (5UQR) (green) oxaloacetate binding site. Oxaloacetate is shown in orange spheres. Most of the amino acid residues forming this site are conserved, except R307 (GltA_Pa_ numbering). D; The left-hand panel shows the open (red) apo-conformation of PrpC_Pa_, the middle panel shows the partially closed (blue) holo-conformation of PrpC_Pa_, and the right hand panel shows a superposition of both conformations of PrpC_Pa_. Note the structural rearrangement in the oxaloacetate-bound PrpC_Pa_ protomer (indicated by the red arrow).

In the PrpC_Pa_ apo-structure (open conformation), each protomer in the asymmetric unit is identical (backbone RMSD of 0.23 Å from a total of 360 Cα atoms). However, in the holo-PrpC_Pa_ structure, the conformation of one of the oxaloacetate-bound protomers (chain D) in the asymmetric unit was different. Each asymmetric unit comprised four monomers of PrpC_Pa_, but only chain D contained an unambiguous electron density for oxaloacetate (Figure 5B). The other chains (chains A, B and C) had an identical conformation to those of apo-PrpC_Pa_. Interestingly, the dimerization partner of chain D, chain C, (shown in orange in Figure 5A) had no oxaloacetate in its active site. This raises the possibility that PrpC_Pa_ exhibits half-of-the-sites reactivity, where only one-half of the identical subunits are active at any given time (47).

GltA_Pa_ has the essential catalytic triad of residues that are also found in the *Sus scrofula* citrate synthase; His265, His306 and Asp363 (Pa numbering). The side-chain orientation in this triad is identical in the majority of apo-PrpC and apo-GltA structures, including PrpC_Pa_ (Figure 5C). In *S. enterica* PrpC, Tyr197 and Leu324 (Tyr184 and Ala311 in PrpC_Pa_) have been proposed to confer substrate specificity (45). The corresponding residues in the citrate synthases are histidine and valine (His227 and Val363 in GltA_Pa_). However, the PrpC from *A. fumigatus* also has histidine and valine in these positions; hence, the precise role(s) of these residues in imparting substrate specificity are still not clear. In addition to binding its substrate, citrate synthase from *E. coli* also binds NADH, and may even be regulated by this compound. The residues important for NADH binding in GltA from *E. coli* are Met112 and Cys206 (44). These residues are also present in GltA_Pa_, but they are absent from PrpC_Pa_. This is consistent with the notion that GltA_Pa_ is probably regulated by NADH (48), whereas this is probably not the case for PrpC_Pa_.

Superposition of the apo-PrpC_Pa_ and holo-PrpC_Pa_ structures highlights the conformational change associated with oxaloacetate binding (Figure 5D). The oxaloacetate- bound PrpC_Pa_ has a more compact configuration, achieved through a 2 Å (average) movement and 7° rotation of the associated domain toward the centre of the dimer. This conformation was also observed in the *Sus scrofa* citrate synthase, where it was described as a “partially closed conformation” (49). The fully closed conformation was observed when both oxaloacetate and Ac-CoA were bound to the enzyme (50). Compellingly, all acyl-CoA bound citrate synthase structures in the PDB contain either bound oxaloacetate or bound citrate. This suggests an ordered reaction sequence. Indeed, in citrate synthase, the binding of oxaloacetate has been biochemically and structurally demonstrated to bring about a conformational change which appears to be critical for the subsequent binding of Ac-CoA (49). Presumably, a similar ordered reaction sequence is associated with PrpC_Pa_. Consistent with the notion that oxaloacetate binding is accompanied by a conformational change in the enzyme, we observed increased thermal stability of PrpC_Pa_ upon the addition of oxaloacetate (Figure S4C).

### Transcriptomics reveals how Pa responds to challenge with exogenous propionate

From the proteomic data, PrpC_Pa_ (and all other enzymes of the 2MCC) were detectable during growth of Pa on succinate as a sole carbon source, so it is possible that the 2MCC also serves a non-canonical, uncharacterised role(s) in Pa physiology. To explore this further, RNA-seq was used to (i) compare the transcriptome of wild-type PAO1 with that of an isogenic Δ*prpC* mutant during growth on succinate, and (ii) examine how the transcriptome is perturbed following exposure of succinate-grown cells to a sub-inhibitory concentration (500 µM) of propionate (added during exponential growth (Figure 6A, S5A- E)).

**Figure 6.**
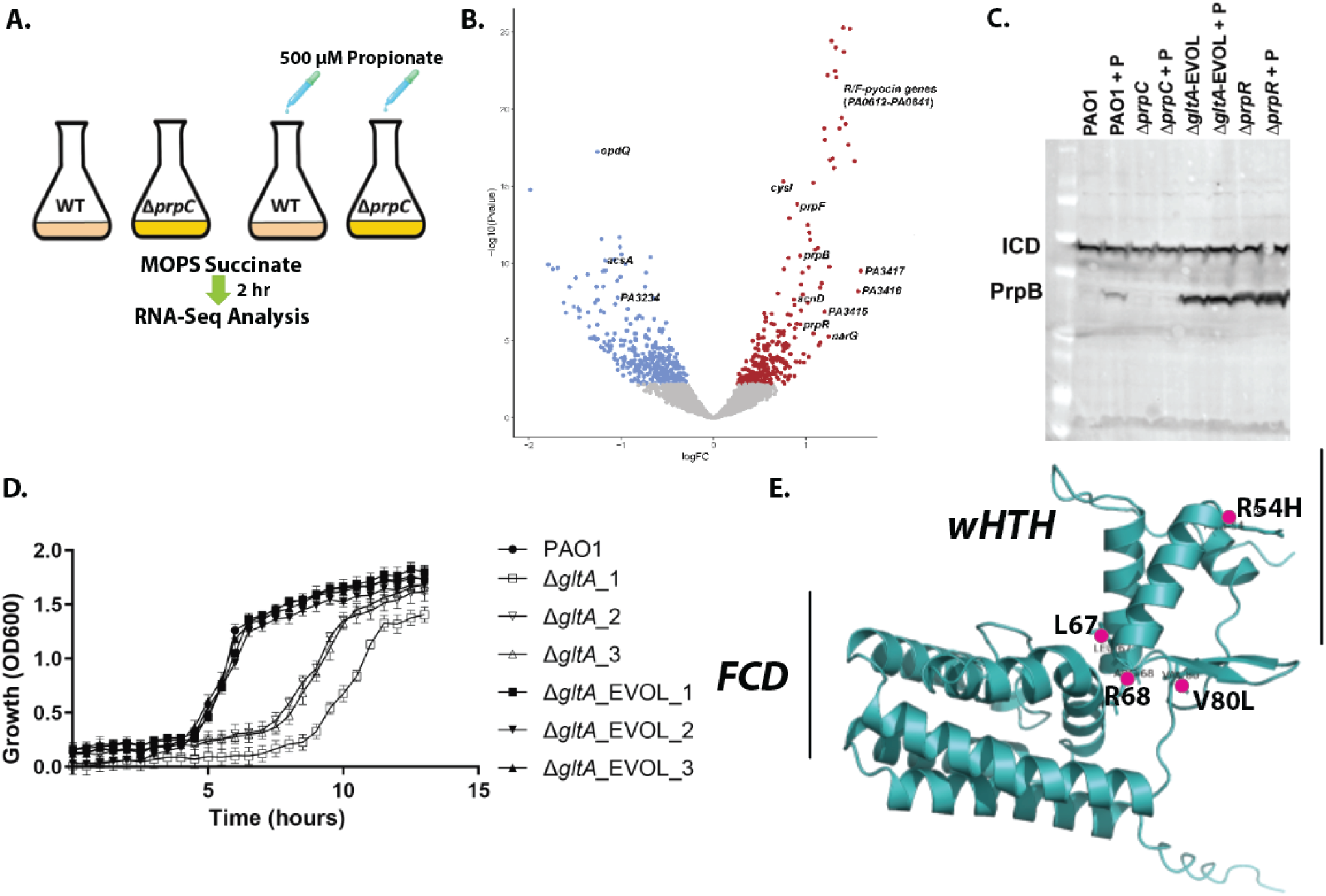
RNA-seq analysis uncovers that propionate exposure induces expression of the *prp* operon and of the genes associated with branched chain amino acid catabolism in Pa. A; Schematic of the experimental design. At OD 0.2, 500 µM sodium propionate was spiked into (triplicate) cultures of PAO1 and the Δ*prpC* mutant. An equal volume of H_2_O was added to the control PAO1 and Δ*prpC* mutant cultures (also grown in triplicate). The cultures were harvested 2 h after the propionate addition, corresponding to an OD_600_ of ≅ 0.6 (exponential growth) and RNA-seq analysis was carried out. B; The panel shows a volcano plot illustrating the log2-fold change in transcript abundance *versus* adjusted *p*-values for wild-type PAO1 grown in MOPS-succinate *versus* wild-type PAO1 grown in MOPS-succinate + 500 µM propionate. Transcripts which are significantly (q- val < 0.05) increased (red) or decreased (blue) in abundance are indicated. Selected transcripts are labelled. C; Western blot showing protein expression levels of PrpB (32.1 kDa) in PAO1, and in the Δ*prpC* mutant, the Δ*gltA*_EVOL mutant, and the Δ*prpR* mutant after exposure to 4 mM propionate (+P) for 3 h. *Iso*citrate dehydrogenase (ICD – 45.6 kDa) served as loading control. Note that the Δ*gltA*_EVOL and Δ*prpR* mutants display constitutively active PrpB expression, independent of propionate addition. Data representative of three independent experiments. D; Growth of Δ*gltA* and Δ*gltA_*EVOL1-3 mutants compared with PAO1 in MOPS-acetate medium. The data are representative of three independent experiments, each performed in triplicate. E; AlphaFold model of PrpR with the locations of the residues mutated and/or deleted in the Δ*gltA_*EVOL_1-3 mutants highlighted. The winged helix-turn-helix (wHTH) motif and the GntR family FadR C-terminal domain (FCD) are shown.

Consistent with the proteomic data, appreciable *prpC* reads were detected in PAO1 during growth on succinate (Figure S5B). This basal expression of the 2MCC enzymes may benefit the cell by priming it ready for rapid propionyl-CoA detoxification/catabolism. Relatively few PAO1 transcripts showed substantial alterations in abundance compared with the Δ*prpC* mutant during growth on MOPS-succinate medium (File S3, Figure S5D). This suggests that the absence of a functional 2MCC does not lead to extensive transcriptional re-programming during *per se*. Transcripts encoding two enzymes (BkdA1, BkdA2) involved in branched chain amino acid (BCAA) catabolism were down-regulated in the Δ*prpC* mutant when compared with the wild-type during growth on succinate. However, this repression was relieved upon exposure of the Δ*prpC* mutant to exogenous propionate (2.2 FC). These data indicate that in the wild-type, flux through the 2MCC during growth on succinate may produce low levels of propionate, and that this impacts on BCAA catabolic gene expression. The source of this propionate could be from the catabolism of endogenously produced, propionyl-CoA-generating amino acids or possibly through reverse operation of the 2MCC. The 2MCC has recently to be shown to be reversible in *M. tuberculosis* to allow optimal metabolism of lactate and pyruvate (51).

The most statistically-significant up-regulated transcripts in the wild-type following challenge with propionate were associated with ORFs PA3415-PA3417 (2.8 FC). These ORFs are predicted to encode a pyruvate dehydrogenase (PDH) or a branched chain amino acid dehydrogenase (File 3, Figure 6B) (52). Immediately adjacent to the PA3415-PA3417 cluster is leucine dehydrogenase (*ldh*, PA3418), required for BCAA catabolism, which was also significantly up-regulated upon propionate exposure. The same ORFs were also up- regulated in the Δ*prpC* mutant after propionate addition, indicating that full catabolism of propionate is not required as a cue to activate the expression of these genes (File S3, Figure S5C). However, given that propionyl-CoA is an intermediate in the metabolism of BCAA, these results likely indicate regulatory cross-talk between expression of the PA3415- PA3417-*ldh* cluster and expression of the enzymes involved in BCAA catabolism. Unexpectedly, *acsA* and PA3233-PA3235 (the ORF cluster that includes the ActP protein) which have putative roles, respectively, in propionate activation and transport, were down- regulated (-2.0 FC) in PAO1 upon propionate addition.

Pa responds to propionate exposure by increasing expression of the *prp* operon (Figure 6B). This up-regulation of the *prp* operon was blocked in the Δ*prpC* mutant (Figure S5E, File S3). We further confirmed the induction of PrpB expression in response to propionate by western blotting (Figure S5F). However, this propionate-induced expression of PrpB was abolished in the Δ*prpC* mutant (Figure S5F). PrpB expression in response to propionate challenge was maintained in a Δ*acsA* mutant, suggesting that propionyl-CoA can also be generated from propionate through alternative routes in Pa. *AcsA* expression is known to be under the control of the response regulator, ErdR, and consistent with this, an Δ*erdR* mutant cannot grown on ethanol or acetate as a sole carbon source (53). Given the dual role that AcsA seems to play in acetate and propionate catabolism, we therefore examined whether a Δ*erdR* mutant also displays aberrant growth on propionate. It did (Figure S5G-I). Surprisingly, the growth deficit of the Δ*erdR* mutant on propionate was even more pronounced than that of a Δ*acsA* mutant. This indicates that additional downstream targets of ErdR, such as ErcS, ErbR or the ethanol oxidation system, may also be required for optimal propionate catabolism in Pa (54).

How is propionate assimilated in Pa and converted to propionyl-CoA, activating the 2MCC operon even when preferable carbon sources are readily available? Carbon catabolite repression (CCR) allows Pa to selectively assimilate a preferred compound when a selection of carbon sources are available. In Pa, CCR is controlled through translational silencing, mediated by Hfq and the small protein Crc (55). Reversing this translational silencing requires the small RNA (sRNA) CrcZ, which sequesters Hfq thereby preventing the latter from binding to target transcripts. CrcZ abundance is controlled by a two-component system, CbrAB, which senses and responds to carbon availability (55). In Pa, *acsA* mRNA harbours a sequence motif located upstream of the *acsA* start codon, which brings acetate assimilation under CCR control. Because they are impaired in CrcZ expression, mutants defective in *cbrB* exhibit a severe growth defect when grown on acetate as a sole carbon source ((56) and Figure S5I). We found that a Δ*cbrB* mutant also had a clear growth defect on propionate (Figure S5H). However, since the Δ*cbrB* mutant maintained inducible PrpB expression upon propionate exposure (Figure S5J), this suggests that CCR does not exert direct control over the 2MCC, but may impact on propionate catabolism indirectly e.g., by affecting *acsA* expression and/or other peripheral targets.

Since CCR did not appear to be directly coordinating the expression of the 2MCC, this prompted us to examine in more detail the role of the GntR-family TF, PrpR (PA0797) in controlling *prp* gene expression. GntR family TFs are typically regulated by ligands that are metabolic substrates/products/cofactors associated with the products of the genes that they regulate (57). Previous studies in *Corynebacterium glutamicum* and *S. enterica* established that 2-methylcitrate (2-MC), the reaction product of PrpC, is a co-activator of the Fis-family PrpR in these species (58, 59). By contrast, PrpR from *M. tuberculosis* is a 4Fe4S protein that employs propionyl-CoA as a co-activator (60). Our observation, that the 2MCC is not induced upon exposure to propionate in a Δ*prpC* mutant (Figure S5E,F), is consistent with 2-MC rather that propionyl-CoA being the co-activator, especially given that propionyl-CoA accumulates in a Δ*prpC* mutant following propionate challenge (Figure 3C).

In contrast to all other species characterized to date, the Δ*prpR* mutant of Pa had no growth defect on any of the carbon sources tested (Figure S5H-J). This suggested that the canonical model of 2MCC regulation by PrpR established for other organisms does not apply in Pa. Remarkably, PrpB was over-expressed in the Δ*prpR* mutant during growth on succinate, independent of the presence or absence of propionate in the medium (Figure 6C). This suggested that Pa PrpR may actually be a *repressor* of the 2MCC, rather than an activator (58). Consistent with this, expression of *prpR* from a plasmid (pUCP20) in the Δ*prpR* mutant was sufficient to repress PrpC expression in this strain (Figure S6A). The predicted PrpR binding motif in Pa, identified by phylogenetic footprinting, is a 12-nucleotide palindrome with the consensus sequence ATTGTCGACAAT (16): this sequence is found upstream of PrpR (84 bp) in PAO1 and PA14. PrpR was recombinantly-expressed and purified to homogeneity (Figure S6B) for electrophoretic mobility shift analyses (EMSA; Figure S6C). These data revealed that PrpR does indeed bind to the upstream region of *prpR*.

Somewhat surprisingly, we found that a Pa citrate synthase mutant (Δ*gltA* – Figure 4A) could also grow on single carbon sources in minimal media, albeit with a prolonged lag phase. Given that PrpC_Pa_ can carry out the condensation of acetyl-CoA with oxaloacetate (Figure 4B) and can therefore potentially substitute for GltA, we suspected that the viability of the Δ*gltA* mutant on single carbon sources might be explained by induction of *prpC*. Consistent with this, and despite repeated attempts at doing so, we were unable to make a Δ*gltA* Δ*prpC* double mutant. If our hypothesis is correct, and given that *prpC* expression is repressed by PrpR, we began to wonder whether *prpR* in the Δ*gltA* mutant might be under strong selection pressure to acquire loss-of-function mutations, thereby boosting PrpC expression. Commensurate with this, successive rounds of sub-culturing of the Δ*gltA* mutant in MOPS-succinate readily yielded heritable derivatives displaying a restoration of rapid growth in this medium. Furthermore, these “evolved” Δ*gltA* mutants also constitutively expressed PrpC and PrpB, independent of propionate addition (Figure 6C, S6A). A similar restoration of rapid growth in MOPS-succinate medium was observed when we deleted *prpR* in the Δ*gltA* background (Figure S6D).

But are loss-of-function mutations in *prpR* the most probable evolutionary path taken by the Δ*gltA* mutant to overcome its metabolic bottleneck? Alternative mechanisms could include mutation of the PrpR binding site upstream of the *prp* operon, duplication of *prpC*, or inactivation of one or more uncharacterized genes involved in modulating PrpR expression. To investigate this further, we made fresh deletions in *gltA* (to minimize the possibility of the strain acquiring additional mutations prior to passaging). Three independent Δ*gltA* mutant colonies were isolated and passaged in MOPS-succinate for two days; all three cultures displayed wild-type levels of growth after this time (Figure 6D, S6D). The three independently evolved Δ*gltA* mutants were sent for whole-genome sequencing alongside the wild-type progenitor (Figure S6E). Strikingly, each of the three evolved Δ*gltA* mutants (EVOL_1-3) had accrued distinct missense mutations in *prpR* giving rise to the amino acid substitutions V80L (EVOL_1) and R54H (EVOL_3) in PrpR, or a 6 bp deletion in *prpR* leading to the loss of amino acids L67 and R68 in PrpR (EVOL_2). These residues were mapped on to the AlphaFold-generated structure of PrpR, which indicated that they fall within and proximal to the conserved winged helix-turn-helix (wHTH) domain of this repressor – a region crucial for the DNA-binding in this family of transcription factors (Figure 6E) (61, 62). Indeed, mutation of residue R52 (equivalent to residue R54 in PqsR) in the *E. coli* MqsR-controlled colanic acid and biofilm regulator (McbR) results in a loss of DNA-binding (57). This provides clear evidence that in the absence of *gltA*, *prpR* is reproducibly mutated to facilitate *prpC* over-expression.

## Discussion

We have carried out a systems-level characterisation of Pa during growth on succinate and propionate as sole carbon sources. This revealed previously undiscovered transcriptional and metabolic crosstalk between several major metabolic pathways/cycles; the 2MCC, BCAA catabolism and the glyoxylate shunt. Our work also provides mechanistic insight into how enzyme promiscuity and regulatory rewiring can rapidly overcome the loss of a key enzyme in the TCA cycle, citrate synthase. We show that Pa can survive the loss of citrate synthase (GltA) through a combination of low, basal-level expression of PrpC, followed by acquisition of loss-of-function mutations in the transcriptional repressor *prpR*. This leads to a compensatory increase in secondary citrate synthase activity through PrpC over-expression (Figure 7A-C).

**Figure 7.**
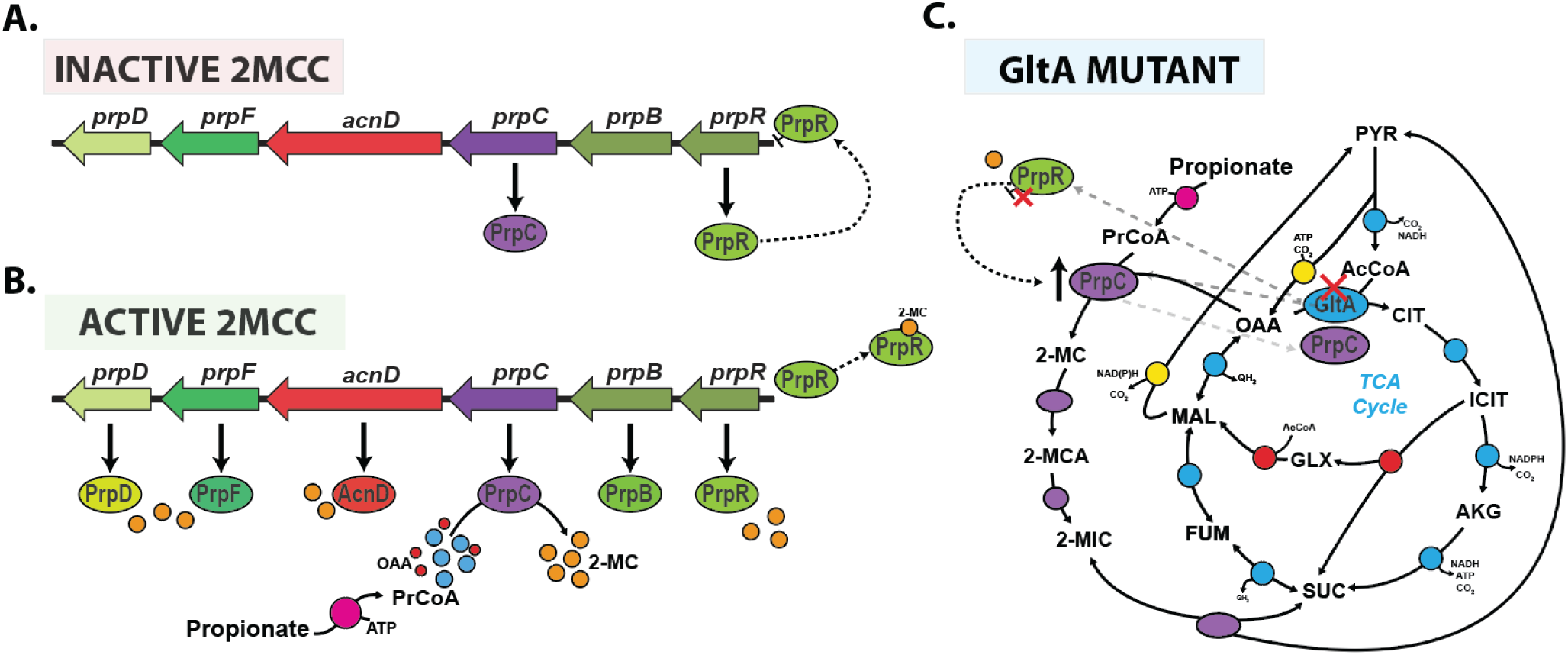
Model for the operation of the 2MCC in Pa. A; During growth in the absence of propionate or propionyl-CoA generating substrates, the 2MCC operon (*prp*) expression is repressed through the binding of PrpR to its upstream promoter region. Incomplete repression of the operon (from basal cellular propionyl-CoA or competing transcriptional activators) results in a basal, low level of *prpC* transcription. B; As the cellular propionyl-CoA levels rise, this metabolite is condensed with oxaloacetate by PrpC, resulting in the formation of 2-MC. 2-MC likely then binds to PrpR, inducing conformational changes which lead to the dissociation of PrpR from the DNA. This de- represses the *prp* operon, allowing expression of the 2MCC enzymes. However, as the concentration of propionyl-CoA falls (due to depletion of propionate or BCAAs due to 2MCC activity) so too does the concentration of 2-MC, which, in turn, leads to re-binding of PrpR to the *prp* promoter region and a resumption in *prp* operon repression. C; In the absence of citrate synthase (GltA), Pa can survive because of the low-level basal expression of PrpC; a promiscuous enzyme that also has citrate synthase activity. However, this low total citrate synthase activity is unable to meet cellular demand, resulting in a severe growth defect and a strong selection pressure to acquire mutations that increase *prpC* expression. Based on our work, it seems that mutations in *prpR* which abolish its repressor activity are the most commonly-selected mechanism for achieving this. These mutations lead to constitutive expression of the *prp* genes, and thus, an increase total cellular citrate synthase activity (compensating for the loss of GltA activity).

We found that Pa responds to propionate exposure by increasing expression of the *prp* operon. This propionate-dependent expression of the 2MCC was unaffected by carbon catabolite repression (CCR) or by deletion of the primary short-chain acyl-CoA synthetase, AcsA. This may reflect the established appetite of Pa for organic acids (1), but it could also be that the primary role of the 2MCC is in propionate detoxification, rather than routine carbon assimilation. It appears that Pa counters rapid propionyl-CoA generation by having an exceptionally responsive 2MCC which promptly degrades inhibitory metabolic intermediates. In *M. tuberculosis*, this detoxification is carried out by the constitutively expressed methylmalonyl-CoA (MMCO) pathway, which can quickly react to sudden changes in propionate concentration and detoxify the cell accordingly. By contrast, the role of the *M. tuberculosis* 2MCC appears to be as a “professional catabolizer”, with a higher overall flux capacity than the MMCO (63). The absence of a functional MMCO in Pa means that this organism depends exclusively on the 2MCC for both the assimilation and detoxification of propionate.

The trade-off between responsiveness to propionate and the accumulation of cytotoxic 2MCC intermediates is a structural weakness of this catabolic arrangement; a weakness that can potentially be exploited to fight Pa infections. Importantly, a synthetic PrpC inhibitor was bacteriostatic against *M. tuberculosis* grown in cholesterol media (cholesterol is broken down by *M. tuberculosis* to yield propionyl-CoA). This suggests that cell-permeable PrpC-specific inhibitors are indeed achievable (64). Considering the structural similarity between PrpC and GltA, it may be possible to generate an inhibitor which targets both enzymes simultaneously. This could be a powerful combination, as a transposon mutagenesis screen indicated that *gltA* is required for the growth of nine different Pa strains from diverse sources when cultured in four infection-relevant growth conditions (LB, M9 glucose, sputum, and serum) (65). Interestingly, most of these strains did not require *gltA* for growth in urine (65, 66). However, the current work highlights the risk of targeting GltA exclusively, since Pa can swiftly compensate for the loss of GltA activity by increasing PrpC expression (through PrpR inactivation) (Figure 7).

Mutations in core metabolic genes are strongly associated with antimicrobial resistance, although our insight the mechanistic basis for this is poorly understood (67–69). Crucially, pathogen lifestyles vary, and this in turn leads to major alterations in the regulatory architecture of primary metabolism. These design variations mean that many of the metabolic innovations that facilitate adaptation to new environments (or to antimicrobial challenge) are pathogen specific.

Can we predict the potential routes of mutation and genetic evolution? Addressing this is a central challenge for evolutionary systems biology, and requires a clear understanding and appreciation of microbial metabolic network diversity. As shown in the current work, large-scale comparative ‘omics analyses, in combination with reverse genetics, can provide mechanistic insights into the complex evolutionary trajectories of underground metabolism. Indeed, the specific path to de-repression of *prpC* expression in Pa (*via prpR* inactivation) that allows the cell to survive in the absence of citrate synthase simply cannot happen in *E. coli* or indeed, in many other human pathogens, due to key differences in metabolic architecture, enzymology, and gene regulation (70). Therefore, the strategic inhibition of organic acid catabolism in Pa through inhibition of PrpC and GltA activity may be a potent mechanism to halt the growth of this pathogen during infection in environments where propionate is abundant.

## Acknowledgements

This work was funded by grants BB/M019411/1 and BB/R005435/1/FAPESP from the BBSRC. SKD was supported by a Herchel Smith Postdoctoral Fellowship. Elements of this work were supported by an EMBO Short Term Fellowship to SKD (7293–2017). CW acknowledges support by the German Federal Ministry for Education and Research (BMBF) through the grants “BioNylon” (FKZ 03V0757) and “LignoValue” (FKZ 031B0344A) and by the German Research Foundation (DFG) through the grant “ePseudomonas” (WI-1796/4-1) within the Priority Programme “eBiotech” (SPP 2240). We thank the Cambridge Centre for Proteomics, including Dr Mike Deery, Mrs Renata Feret and Prof Kathryn Lilley for proteomics support.

## Competing Interests Statement

The authors declare that they have no competing interests.

## Materials and Methods

### Growth conditions

Unless otherwise indicated, *P. aeruginosa* strain PAO1 (71) was routinely grown in lysogeny broth (LB LENNOX) (Oxoid Ltd) at 37°C with good aeration (shaking at 250 rpm). The strains used in this study are listed in Table S1A. The overnight pre-cultures were started from separate clonal source colonies on streaked LB agar plates. Strains were cultured in MOPS (morpholinepropanesulfonic acid) media with the relevant carbon sources (72). Cell growth was monitored as optical density in a spectrophotometer (BioSpectrometer®, Eppendorf) at a wavelength of 600 nm (OD_600_). A previously determined conversion factor of 0.42 g cell dry weight (CDW) per OD_600_ unit was used to calculate biomass specific rates and yields from the obtained OD_600_ values (73).

### Transcriptomics (RNA-Seq)

*P. aeruginosa* strain PAO1 and Δ*prpC* were grown in 40 ml MOPS with succinate (30 mM) as the sole carbon source (six flasks per strain) at 37°C with good aeration (shaking at 250 rpm) in baffled flasks (500 ml volume). At OD_600_ 0.2, 500 µM sodium propionate was spiked into three of the PAO1 cultures and three of the Δ*prpC* cultures. An equal volume of H_2_O was added to the control PAO1 and Δ*prpC* cultures. After 2 hr, an aliquot (5 ml) of culture was removed from each sample. At this stage, the culture OD_600_ was ≅ 0.7 (exponential growth). These aliquots were added to an equal volume of RNAlater® RNA stabilization solution. RNA was then isolated using a RNeasy Mini Kit (Qiagen). Ribosomal RNA (rRNA) was subsequently depleted from each RNA sample (5 µg each) using the bacterial Ribo-Zero rRNA Removal Kit (Illumina). The integrity of the RNA was evaluated using an RNA 6000 Nano LabChip and an Agilent 2100 Bioanalyzer (Agilent Technologies, Germany). Twelve indexed, strand-specific cDNA libraries were prepared, and samples were sequenced on an Illumina HiSeq 2000 with a 51 bp single-end read length (GATC Biotech, Germany). The sequencing data are deposited at ArrayExpress (accession number E-MTAB-10077).

### Reads mapping and annotations

The FASTQ files were mapped to the PAO1 genome obtained from the Pseudomonas Genome Database (PGD) (http://www.pseudomonas.com/) using Bowtie v.0.12.8 [38]. The sequence reads were adaptor clipped and quality trimmed with trimmomatic (74) using the default parameters. The Integrative Genomics Viewer was used to visually inspect mapping quality and the absence of *prpC* reads in the Δ*prpC* mutant. Read summarization was performed using featureCounts (75). DESeq2 was employed to analyse differentially-expressed genes (76). Annotations of differentially expressed genes were obtained from the reference annotation of the Pseudomonas genome available at the PGD website. Genes were considered as significantly induced or repressed when their adjusted-P value was <0.05 (File S3).

### Quantitative proteomic analysis

*P. aeruginosa* PAO1 cells (OD_600_ = 0.5, 30 ml) were grown at 37°C in 40 ml MOPS with succinate (30 mM) or propionate (40 mM) as the sole carbon source, with good aeration (shaking at 250 rpm) in baffled flasks (500 ml volume). Cultures were grown and analysed in triplicate. The cell pellets were resuspended in 2 ml lysis buffer (100 mM Tris-HCl, 50 mM NaCl, 10% (v/v) glycerol, 1 mM *tris*(2-carboxyethyl)phosphine (TCEP), pH 7.5) containing one cOmplete Mini protease inhibitor cocktail (Roche). Following three rounds of sonication (3 × 10 sec) on ice, supernatants were clarified by sedimentation (21130 × *g*, 15 min, 4°C) in an Eppendorf 5424R centrifuge. Aliquots (100 μg) of each sample were reduced with TCEP, alkylated with iodoacetamide and labelled with Tandem Mass Tags (TMTs). TMT labelling was carried out according to the manufacturer’s protocol.

### LC-MS/MS

LC-MS/MS analyses were carried out using a Dionex Ultimate 3000 RSLC nanoUPLC (Thermo Fisher Scientific Inc, Waltham, MA, USA) system in-line with a Lumos Orbitrap mass spectrometer (Thermo Fisher Scientific Inc, Waltham, MA, USA) (27). Separation of peptides was performed by C18 reverse-phase chromatography at a flow rate of 300 nL/min using a Thermo Scientific reverse-phase nano Easy-spray column (Thermo Scientific PepMap C18, 2 µm particle size, 100 Å pore size, 75 µm i.d. x 50 cm length).

### Proteomic data analysis

Proteome Discoverer v2.1 (Thermo Fisher Scientific) and Mascot (Matrix Science) v2.6 were used to process raw data files. The data were aligned with the UniProt *Pseudomonas aeruginosa* (5584 sequences) common repository of adventitious proteins (cRAP) v1.0. The R package, MSnbase (77), was used for processing proteomics data. Protein differential abundance was evaluated using the Limma package (78). Differences in protein abundances were statistically determined using Student’s *t*-test with variances moderated by Limma’s empirical Bayes method. *P*-values were adjusted for multiple testing by the Benjamini- Hochberg method (79). Proteins were considered as increased or decreased in abundance, when their log_2_ fold-change was >1 or <−1, respectively, and their *p*-value was <0.05. The mass spectrometry proteomics data have been deposited to the ProteomeXchange Consortium *via* the PRIDE (55) partner repository with the data set identifier PXD015792.

### Genome sequencing

Genomic DNA was extracted from PAO1 and three evolved Δ*gltA* mutants (EVOL_1-3) using a GeneJET Genomic DNA Purification Kit following 50 generations of growth in MOPS- succinate. Genome sequencing of all four strains was carried out by MicrobesNG (http://www.microbesng.uk), and the reads were analysed and displayed using IGV (80).

### Construction of in-frame *P. aeruginosa* PAO1 deletion mutants

Flanking regions 800-1000 bp upstream and downstream of the desired genes were PCR amplified. The upstream and downstream regions were then overlapped and cloned into the suicide vector, pEX19Gm, using Gibson assembly as described previously (81). The resulting deletion plasmid was then introduced into *P. aeruginosa* by electroporation and selected for on LB plates containing 50 µg/ml gentamicin. Deletion mutants were identified *via* SacB-mediated sucrose counter-selection and confirmed by PCR. Primers used are described in Table S1B.

### Construction of luciferase reporter strains

Transcriptional reporter constructs were made by fusing the upstream promoter sequences of the indicated genes with the *luxCDABE* cluster using the primers listed in Table S1B. The purified PCR products were digested and directionally ligated into the multiple cloning site of plasmid pUC18T-mini-Tn7T-lux-Gm (82). The mini-Tn7-lux element was introduced into PAO1 (where it integrated into the chromosome) by electroporation along with the helper plasmid pTNS2, as previously described (83). Luciferase and OD_600_ readings were measured using a BMG Labtech FLUOstar Omega microplate reader. Strains were cultured in MOPS media with the indicated carbon sources (100 µL) in 96-well microplates (Greiner bio-one, F- Bottom, Black) covered with gas permeable imaging seals (4titude - 4ti-0516/96). Luciferase expression was assessed during exponential growth. Growth was measured by taking OD_600_ readings simultaneously with the luminescence readings. Luciferase readings were expressed as relative luminometer units (RLU) normalised to OD_600_ in order to control for growth rate differences in the selected carbon sources.

### ^13^C fluxomics

Starter cultures were prepared by inoculating LB medium with a loop of freshly plated PAO1. After 6 hr of incubation, 50 µL of cell suspension was transferred to a second culture of MOPS minimal medium containing the desired substrate (see below). Subsequently, exponentially growing cells were used as an inoculum for the main cultures. In the main cultures, PAO1 was grown in 25 mL of minimal medium in baffled shake flasks (250 mL volume) with good aeration (shaking at 200 rpm, 37°C) in an orbital shaker (Aquatron, Infors AG, Switzerland). In these conditions, the oxygen level is maintained above 80% of saturation (73).

For the second and main cultures, PAO1 was grown in MOPS minimal media with 40 mM propionate or 30 mM succinate as the sole carbon source (i.e., 120 mM carbon in each case). For ^13^C flux experiments, naturally labelled propionate and succinate was replaced with separate tracers (three for propionate, two for succinate) to maximise dataset resolution and to accurately determine substrate uptake. Naturally labelled propionate was substituted with [1,3-^13^C_2_] sodium propionate (99%), [3-^13^C] sodium propionate (99%) and an equimolar mixture of [U-^13^C] sodium propionate (99%) and naturally labelled sodium propionate (Sigma-Aldrich, Poole, Dorset, UK). Naturally labelled succinate was substituted with 99% [1,4-^13^C_2_] sodium succinate, 99% [2,3-^13^C] sodium succinate or an equimolar molar 1:1 mixture of [U-^13^C] sodium succinate (obtained from Cambridge Isotope Laboratories, Inc., Andover, MA, USA) and naturally labelled sodium succinate.

In cultures incubated with ^13^C-tracer, the inoculum (initial OD < 0.02) was always kept below 1% of the final sampled cell concentration to exclude potential interference of non-labelled cells on subsequent calculation of flux (84). Mass isotopomer labelling analysis of proteinogenic amino acids, mass isotopomer labelling analysis of cell sugar monomers (glucose, ribose and glucosamine), metabolic reaction network and flux calculation were carried out as described in (20).

### Quantification of substrates and products

Propionate and succinate, as well as organic acids (citric acid, α-ketoglutaric acid, gluconic acid, 2-ketogluconic acid, pyruvic acid, succinic acid, lactic acid, formic acid, fumaric acid, and acetic acid) were quantified in filtered culture supernatants (Costar® Spin-X® 0.22 μm) using isocratic high-performance liquid chromatography (Agilent 1260 Infinity series, s HPX- 87H column operating at 65°C and a flow rate of 0.5 mL min^-1^) equipped with refractive index (RI) and ultraviolet (UV) detectors (210 nm) with 12 - 50 mM H_2_SO_4_ as an eluent (85). Concentrations were determined from commercial standards which were analysed on the same run. These data were then used to calculate specific uptake and formation rates, and biomass yields for propionate, succinate, and secreted by-products, respectively (File S2).

### Calculation of redox cofactor and ATP balances

Total production of reduced cofactors was determined by summing up all cofactor-forming fluxes taking into account substrate-dependent cofactor specificities (86–88). Anabolic NADPH requirements and anabolically-produced NADH were estimated from the biomass composition (73, 89) and measured specific growth rates. Surplus NADPH was considered to be converted into NADH *via* the activities of a soluble (SthA, PA2991) and a membrane- bound, proton-translocating (PntAB, PA0195-PA0196) pyridine nucleotide transhydrogenases (34).

ATP: The total ATP demand was calculated by summing up (i) the anabolic demand needed for biomass building block synthesis, and (ii) polymerisation estimated from cell composition multiplied with the corresponding specific growth rate on each substrate (20, 73). We also took into account the costs of growth-associated maintenance (GAM) and non-growth-associated maintenance (NGAM) (90) and ATP costs for substrate activation; the full reaction reference network is shown in File S2, (91). The ATP synthesized by oxidative phosphorylation *via* the respiratory chain was estimated assuming a P/O ratio of 1.875 for NADH and PQQH_2_ (92) and 1.0 for FADH_2_ and other quinone (QH_2_) carriers (93), respectively. The anabolic ATP requirement was calculated from published biomass composition data for pseudomonads (mainly protein, RNA and lipid synthesis) inclusive of the costs of polymerising the precursors of these components (73, 94). The GAM and NGAM costs for pseudomonads was previously modelled using genome-scale models (90, 95). Here, an ATP surplus represents the amount of ATP available to fulfil remaining cellular ATP-consuming tasks.

### Western-blot analysis

Equal amounts of protein (10 µg) were resolved on a 12% SDS—polyacrylamide gel. The resolved proteins were blotted onto a nitrocellulose membrane, which was blocked with 5% (w/v) skimmed milk power in TBS buffer. The membranes were probed with rabbit-derived anti-ICD antibodies, anti-PrpC antibodies or anti-PrpB antibodies (polyclonal antibodies raised by BioGenes.De). Following washing to remove excess primary antibody, the membranes were then probed with IRDye® 800CW goat anti-rabbit IgG secondary antibodies (926–32211). Bands were visualized on an Odyssey Infrared Imaging System (LI-COR Biosciences).

### Enzymatic assays

The 2-methylcitrate synthase (2-MCS) activity of PrpC was measured using a method described by Srere *et al* (96) except that propionyl-CoA (PrCoA) was used instead of acetyl- CoA (AcCoA). Briefly, the condensation reaction of oxaloacetate (OAA) and PrCoA facilitated by PrpC generates free coenzyme A (CoA). The free CoA thiol group on the liberated CoA reacts with 5,5’-dithio*bis*(2-nitrobenzoic acid) (DTNB) to yield 2-nitro-5-thiobenzoate (TNB^2-^) anions. TNB^2-^ is coloured and its formation can be monitored at 412 nm. The initial rate was calculated from the rate of change of the A_412_ assuming an extinction coefficient for TNB^2-^ of 14,150 M^-1^ cm^-1^. The reaction mixtures contained buffer (50 mM HEPES pH 7.5, 0.1 M KCl, and 0.54 M glycerol), substrates (OAA and PrCoA at the indicated concentrations) and 0.15 mM DTNB. Reaction mixtures were equilibrated at 37°C for 5 min before the reaction was initiated by the addition of PrpC (to a final concentration of 240 nM). The reaction was kept in 37°C and the A_412_ was measured in a BioSpectrometer® (Eppendorf). Kinetic parameters were calculated using best-fit nonlinear regression and plotted using GraphPad Prism version 6. The citrate synthase (CS) activity of PrpC was measured using the method above but AcCoA in place of PrCoA. The 2-MCS and CS activity of GltA was measured using PrCoA and AcCoA respectively.

### Protein expression

The PCR-amplified ORFs of *prpC*, *gltA* and *prpR* were cloned into the expression vector pET- 19m, which introduces a TEV-cleavable N-terminal hexahistidine tag onto each protein. For purification of the His_6_-tagged proteins, the cells were grown in LB medium (1 L) at 37°C with good aeration to *A*_600_ = 0.5. The temperature was then lowered to 16°C, and isopropyl 1-thio-β-d-galactopyranoside was added to 1 mM final concentration to induce expression of the cloned genes. The induced cultures were grown for a further 16 h and then harvested by sedimentation (6000 × *g*, 4°C, 15 min). The cell pellet was resuspended in 20 ml of buffer A (50 mM sodium phosphate, 100 mM NaCl, 10% (v/v) glycerol, pH 8.0, containing one dissolved cOmplete™ EDTA-free Protease Inhibitor cocktail tablet (Roche)), and the cells were ruptured by sonication (3 × 10 s, Soniprep 150, maximum power output). The cell lysate was clarified by centrifugation (11,000 × *g*, 4 °C, 30 min), and the supernatant was filtered through a 0.45-μm filter. The filtered lysate was then loaded onto a 5 ml Ni-NTA Superflow column (Qiagen), and the column was washed with buffer A containing 10 mM imidazole. The His_6_-tagged proteins were eluted with buffer A containing 250 mM imidazole. His_6_-tagged TEV-protease (1 mg) was added to the purified protein solution, and the mixture was dialyzed overnight at 4°C against 2 litres of buffer B (20 mM Tris-HCl, 50 mM NaCl, 5% (v/v) glycerol, pH 7.5). Uncleaved protein and the His_6_-TEV protease were removed by batch extraction in a slurry of Ni-NTA resin equilibrated in buffer B. The unbound (cleaved) protein was concentrated to the desired concentration using an Amicon® Ultra-4 Centrifugal Filter (10 kDa NMW cut-off).

### Protein crystallization

PrpC. Crystallization conditions were screened using the sitting drop vapour diffusion technique with approximately 13-15 mg/mL of purified PrpC solution. Optimization conditions were prepared using dragonfly® discovery system (TTP LabTech). Protein drops were generated using an automated nanoliter liquid handler mosquito® HTS (TTP LabTech). PrpC apo- and holo- (OAA-bound) crystals were obtained in a 1:1 ratio of protein and reservoir solution (100-200 mM bis-Tris pH 5.5, 20-30% (w/v) PEG 3350, and 0.1% d-xylose). To obtain the OAA-bound structure of PrpC, the crystallization solution additionally contained 2.5 mM oxaloacetate. All crystals were grown for 2-11 days at 19°C, and were cryoprotected with 25% (v/v) glycerol and 75% (v/v) reservoir solution prior to mounting in nylon loops (Hampton Research). Mounted crystals were flash frozen in liquid nitrogen prior to data collection.

GltA. Purified GltA at a concentration of 20-25 mg/mL was crystalized by sitting drop vapour diffusion. Crystals were grown for 7 days at 19°C, and were cryoprotected with 25% (v/v) glycerol prior to being mounted and flash frozen for data collection.

### X-ray diffraction, structure determination and refinement

PrpC. Diffraction data was collected on beamline MX-I03 at the Diamond Light Source Synchrotron (DLS, Didcot, UK). The parameters for the data collection were as follows; Omega (Ω) Start: 62.0°, Ω Oscillation: 0.10°, Total Oscillation: 180°, Total Images: 1800, Exposure Time: 0.050 s. Diffraction images were processed using Xia2 DIALS (97). The structure was determined by molecular replacement using Phaser (98) with the atomic coordinates of the PrpC from *Coxiella burnetii* (PDB entry: 3TQG) as the search model. Automated refinement was performed using Refmac5 (99) and PHENIX.refine (100). Manual modelling and refinement were performed in COOT (101). Data collection and refinement statistics are listed in Table S1D.

GltA. Data was collected on MX-I03 beamline at the Diamond Light Source Synchrotron (DLS, Didcot, UK). The parameters for the data collection were as follows; Omega (Ω) Start: 0°, Ω Oscillation: 0.20°, Total Oscillation: 240°, Total Images: 1200, Exposure Time: 0.050s. Diffraction images were processed using Xia2 DIALS (102). The structure was determined by molecular replacement using Phaser (98) with the atomic coordinates of the type II citrate synthase from *Vibrio vulnificus* (PDB entry: 4E6Y) as the search model. Automated refinement was performed using Refmac5 (103) and PHENIX.refine (104). Manual modelling and refinement were performed in COOT (105). Data collection and refinement statistics are listed in Table S1D.

### Analytical ultracentrifugation

Analytical ultracentrifugation-sedimentation velocity (AUC-SV) was conducted in the Department of Biochemistry (University of Cambridge) Biophysics Facility. Samples were dialyzed overnight at 4°C against a buffer solution containing 100 mM NaCl and 50 mM Tris-HCl pH 7.5 to remove traces of glycerol. Data were collected using an An60Ti analytical rotor (Beckman Coulter) in a Beckman Optima XL-I ultracentrifuge with absorbance and interference optical detection systems. Protein solution (400 μL volume, concentration approximately 1 mg mL^-1^) and the reference solution (dialysate) were added to the Epon (epoxy) double-sector centrepieces. All samples were sedimented at 40,000 rpm and 20°C. Absorbance data (A_280_) were collected in intervals of 2 min and interference scans were taken every 1 min. The viscosity and density of the buffer used in the experiments were estimated with SEDNTERP. Data analysis were conducted using SEDFIT.

### Protein thermal stability

Differential scanning fluorimetry experiments were carried out using a CFX Connect RT-PCR detection system (BioRad) and using Hard-Shell® 96-well PCR plates (BioRad) which are compatible with the excitation and emission wavelength of SYPRO orange. The temperature range was 4-95°C with an increment of 1°C every 45 s. The fluorescence was measured every 15 s. Data was analysed using GraphPad Prism 6.

### EMSA analysis

The region upstream of *prpR* (250 bp, including the 12-nucleotide palindrome) was PCR- amplified. The forward primer contained a 6-FAM (6-carboxyfluorescein) tag (Sigma). EMSA reaction mixtures (25 μL volume) contained buffer (20% (v/v) glycerol, 30 mM Tris-HCl (pH 8.0), 1 mM MnCl_2_, 120 mM KCl, 1 mM MgCl_2_) supplemented with 5 pM 6-FAM-labeled probe, up to 2 µM recombinant His_6_-PrpR, 240 µg/mL bovine serum albumin (BSA), and 15.2 µg/mL poly[d(I-C)]. After incubation at 21°C for 60 min, individual samples were applied to a 6% polyacrylamide gel (Novex) prepared in Tris-borate-EDTA buffer. The samples were electrophoresed in the same buffer system for 45 min at 120 V. The gels were then imaged using an Odyssey imager (Li-Cor Biosciences). In the competition EMSA, unlabelled competitor probes harbouring specific nucleotide substitutions were added in 50-fold molar excess relative to the labelled probes.

### LC-MS analysis of propionyl-CoA

Sampling, analysis and quantification of propionyl-CoA and other CoA esters was carried out as described previously (106). Briefly, cells from 8 mL cultures grown to OD_600_ 2 were pelleted and re-suspended in 200 µL “supercool” ultra-pure water (0°C) and 1 mL quenching-extraction buffer (95% acetonitrile, 25 mM formic acid, -20°C). The mixture was vortexed then kept on ice for 10 min, and finally, centrifuged (3 min, 0°C). The supernatants were transferred into 3 mL of ultra-pure water, then snap-frozen in liquid nitrogen and lyophilized (Alpha 3-4 LSCbasic, Christ, Germany). The freeze-dried samples were diluted in 500 µL pre-cooled resuspension buffer (25 mM ammonium formate, pH 3.0, 2% methanol, 4°C) and immediately analysed by LC-MS (a QTRAP 6500+ (AB Sciex, Darmstadt, Germany) coupled to an HPLC system (Agilent Infinity 1290)). Commercial standards were used for quantification. Final concentrations are given as nmol per gram dry cell weight (DCW).

## Notes

### Competing Interest Statement

The authors have declared no competing interest.

