## Supplementary_Figures for "Systems-wide dissection of organic acid assimilation in *Pseudomonas aeruginosa* reveals a novel path to underground metabolism"

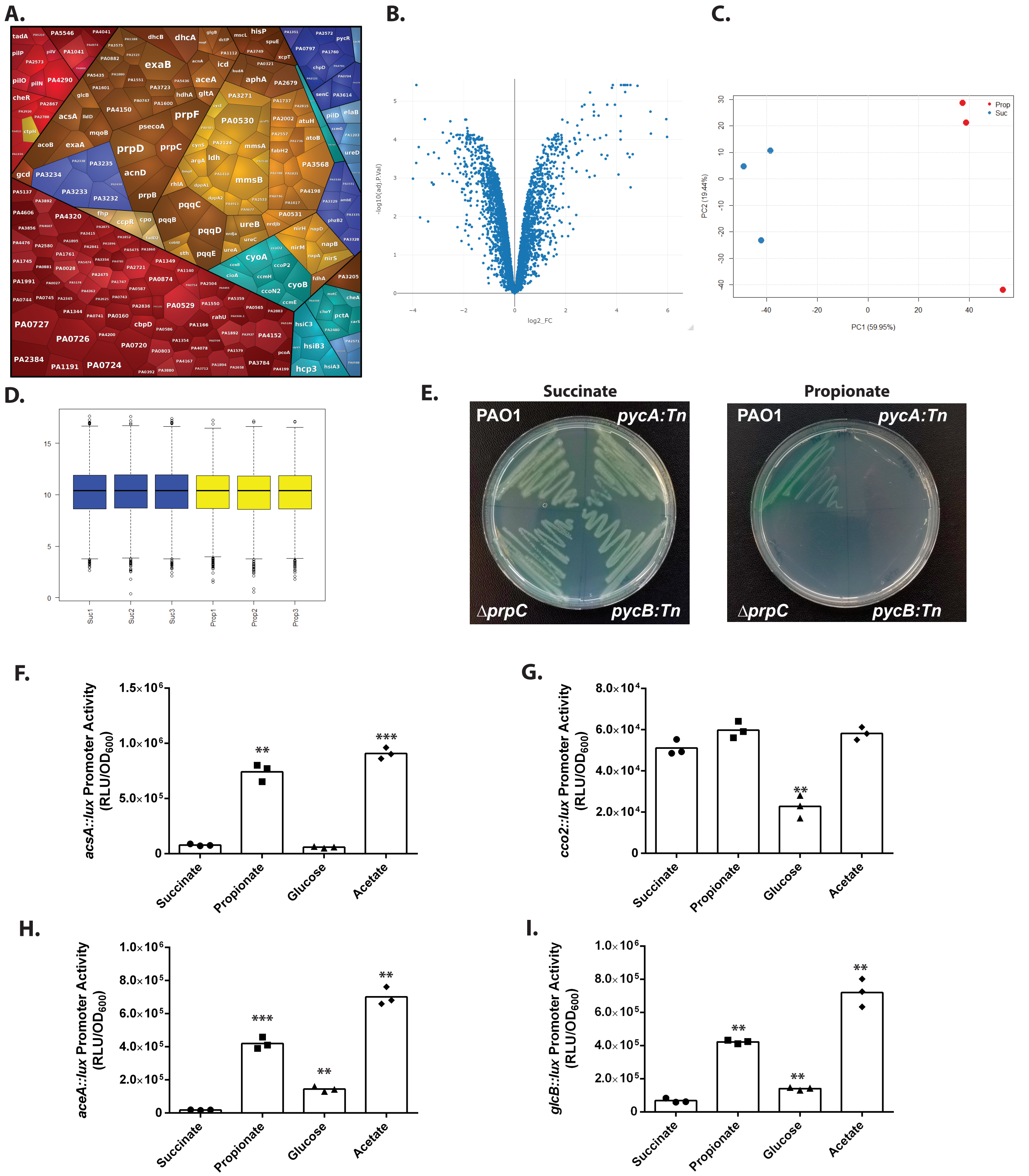


**Figure S1; Proteomic analysis of Pa cultured in MOPS-succinate *versus* MOPS-propionate.**

**A;** A Voronoi tessellation illustrating the proteins which are increased in abundance during exponential growth of PAO1 in MOPS-propionate compared with growth in MOPS-succinate. Notable proteins include those in the ORF cluster (PA3232-PA3235, blue) containing the putative ActP transporter protein (PA3234), enzymes in the 2MCC, glyoxylate shunt and TCA cycle (brown), cytochrome oxidase components (turquoise) and enzymes involved in branched chain amino acid catabolism (tan).

**B;** Volcano plot illustrating $\log_2$ fold change in Pa protein abundance versus adjusted p-values for the MOPS-propionate versus MOPS-succinate datasets.

**C;** Principal component analysis (PCA) of the proteomic data from the Pa grown in MOPS-propionate (red) and MOPS-succinate (blue).

**D;** Box-and-whisker plot illustrating the normalised MOPS-propionate (yellow) and MOPS-succinate (blue) replicates.

**E;** Pa pyruvate carboxylase (*pycA*::Tn and *pycB*::Tn) transposon mutants cultured on MOPS-succinate agar or MOPS-propionate agar (as indicated) alongside PAO1 and Δ*prpC*. Note that the pyruvate carboxylase mutants cannot grow on propionate as a sole carbon source. The plates were photographed after 24 h incubation. The data are representative of two independent experiments performed in triplicate.

**F-I;** Luciferase activity in *P. aeruginosa* PAO1 carrying chromosomal *promoter::lux* fusions for the indicated promoters, cultured in MOPS-succinate, MOPS-propionate, MOPS-glucose, or MOPS-acetate. **F;** *acsA*, **G;** *cco2*, **H;** *aceA*, **I;** *glcB*. Values normalised to OD600 (RLU/OD600). The data represent three biological replicates per sample. The data were analysed using GraphPad Prism (V 6.01) and statistical significance was determined with an unpaired parametric t-test with Welch's correction. Statistically significant differences in RLU/OD600 are indicated as follows ns: $p > 0.05$, * $p \leq 0.05$, ** $p \leq 0.01$, *** $p \leq 0.001$.

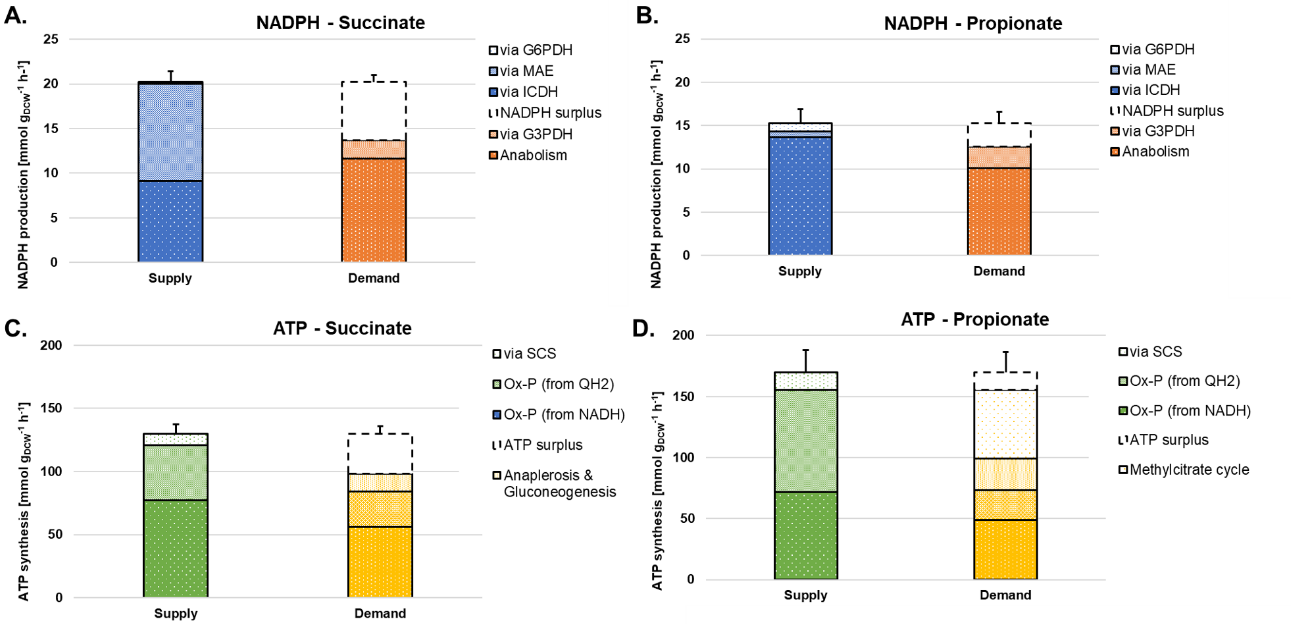


**Figure S2; Quantitative analysis of redox and energy supply and demand for Pa grown on succinate (A, C) or propionate (B, D) as a sole carbon source.** Reactions linked to NADPH (A, B) and ATP (C, D) metabolism were calculated from the obtained fluxes (Figure 2). Values are given as absolute fluxes (mmol $g^{-1}$ $h^{-1}$) and are related to the specific carbon uptake rate (see File S2). G6PDH; glucose 6-phosphate dehydroge-nase, MAE; malic enzyme, ICDH; isocitrate dehydrogenase(s), G3PDH; glyceraldehyde 3-phosphate dehy-drogenase, SCS; succinyl-CoA synthase, Ox-P; oxidative phosphorylation.

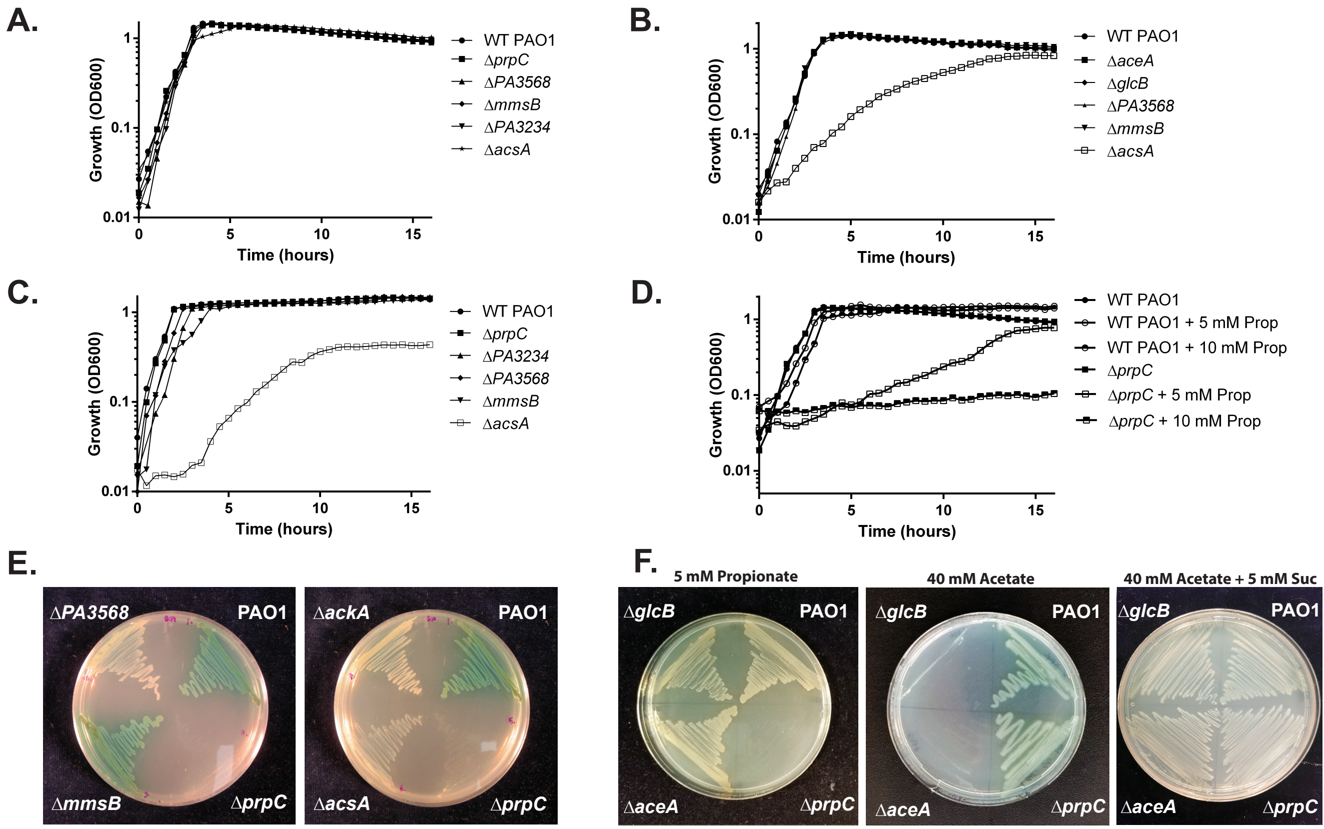


**Figure S3; Growth characteristics of selected mutants in MOPS-succinate, MOPS-propionate, and MOPS-branched chain amino acids media.**

**A;** Growth curves of PAO1, and the Δ*prpC*, ΔPA3568, Δ*mmsB*, ΔPA3234 and Δ*acsA* mutants cultured in MOPS-succinate. Data representative of three independent experiments performed in triplicate.

**B;** Growth curves of PAO1, and the Δ*aceA*, Δ*glcB*, ΔPA3568, Δ*mmsB* and Δ*acsA* mutants cultured in MOPS-propionate. Data representative of three independent experiments performed in triplicate.

**C;** Growth curves of PAO1, and the Δ*prpC*, ΔPA3234, ΔPA3568, Δ*mmsB* and Δ*acsA* mutants cultured in MOPS-acetate. Data representative of three independent experiments performed in triplicate.

**D;** Growth curves of PAO1, and the Δ*prpC* and ΔPA3234 mutants cultured in MOPS-succinate containing 0 mM, 5 mM and 10 mM propionate. Data representative of three independent experiments performed in triplicate.

**E;** Growth of PAO1, and the Δ*prpC*, ΔPA3568, Δ*mmsB*, ΔPA3234, Δ*acsA* and Δ*ackA* mutants cultured on MOPS + branched chain amino acids (BCAA) as a sole carbon source (2 mM each of L-isoleucine, L-valine, and L-leucine). The plates were photographed after 48 h. Data representative of two independent experiments performed in triplicate.

**F;** Growth of PAO1, and the Δ*aceA*, Δ*glcB* and Δ*prpC* mutants cultured on 5 mM MOPS propionate, 40 mM acetate and 40 mM MOPS-acetate + 5 mM succinate. The plates were photographed after 48 h. The data are representative of two independent experiments performed in triplicate.

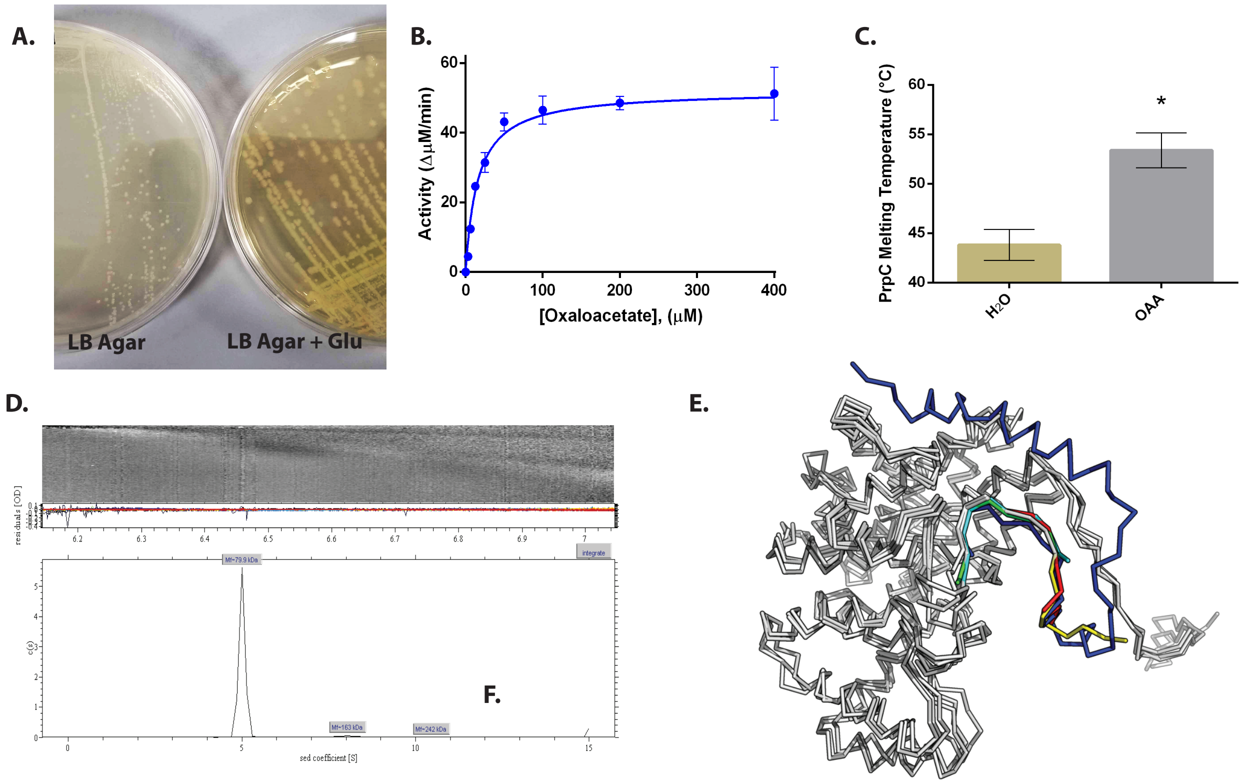


**Figure S4; Supplementary data for structural analysis.**

**A;** The growth defect leading to a small colony size in the Δ*gltA* mutant is partially complemented by supplementing the LB-agar with additional glutamate (5 mM). The data represent two independent experiments, each performed in triplicate.

**B;** Oxaloacetate-dependence of the 2-methylcitrate synthase activity of $PrpC_{Pa}$. The concentration of propionyl-CoA was fixed at 0.5 mM. The data represent two independent experiments, each performed in triplicate.

**C;** Oxaloacetate (200 μM) binding to $PrpC_{Pa}$ leads to a 10°C increase in the melting temperature of the enzyme ($P < 0.0001$, two-tailed unpaired student t-test). The data represent two independent experiments, each performed in triplicate.

**D;** Determination of $PrpC_{Pa}$ (1 mg mL-1) oligomeric state by analytical ultra-centrifugation sedimentation velocity. One hundred and twenty-eight absorbance scans were recorded at 280 nm. Given that the theoretical molecular mass of a $PrpC_{Pa}$ monomer is 41.7 kDa, the single peak of 79.9 kDa suggests that $PrpC_{Pa}$ likely forms a dimer in solution. The calculated frictional coefficient of 1.2 indicates that $PrpC_{Pa}$ is likely globular.

**E;** Alignment of secondary structural elements in a $PrpC_{Pa}$ protomer against other PrpC structures available on the PDB - *Coxiella burnetii*, *S. enterica* (*Typhimurium*), and *Mycobacterium tuberculosis* (PDB entry: 3TQG, 3O8J, and 3HWK). Most of the structures have moderate to high amino acid sequence identity (40-60%) with $PrpC_{Pa}$. The structure similarity analysis tool, PDBeFold, revealed 95% secondary structural identity with an RMSD of 1.5 Å, indicating that PrpC is structurally conserved across these bacterial species. Eukaryotic PrpC (from *Aspergillus fumigatus,* sharing only 24% sequence identity with $PrpC_{Pa}$) has an additional 50 amino acids residues which forms an extra loop and helix at the N-terminus.

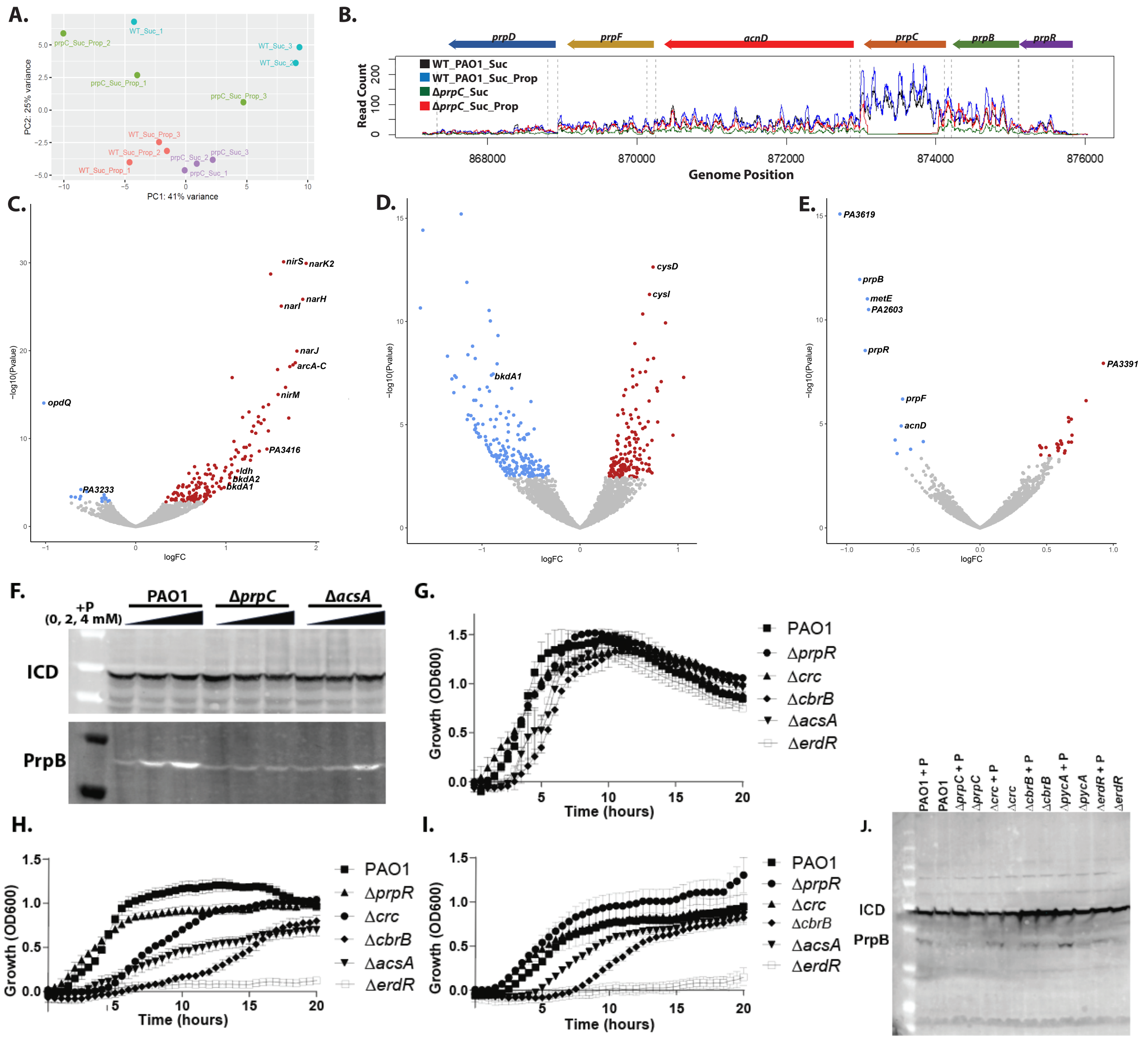


**Figure S5; Transcriptomic analysis (DESeq2) of Pa wild-type and the Δ*prpC* mutant grown on MOPS-succinate versus MOPS-succinate containing 500 µM propionate for 2 h.**
**A;** A principal component(s) analysis (PCA) scores plot of the RNA-seq replicates (triplicates). Wild-type cultured in MOPS-succinate (blue), wild-type cultured in MOPS-succinate + propionate (red), Δ*prpC* mutant cultured in MOPS-succinate (purple), Δ*prpC* mutant cultured in MOPS-succinate + propionate (green).
**B;** The panel shows the sequencing (transcript) reads corresponding to the 2MCC operon for PAO1 and the Δ*prpC* mutant ± propionate. The prpC transcript reads are absent from the Δ*prpC* mutant with no obvious polar effects on the downstream ORFs.
**C;** The panel shows a volcano plot illustrating the $\log_2$-fold change in transcript abundance versus adjusted p-values for the Δ*prpC* mutant grown in MOPS-succinate versus the same mutant grown in MOPS-succinate + 500 µM propionate. Transcripts which are significantly (q-value < 0.05) increased (red) or decreased (blue) in abundance are indicated. Selected transcripts are labelled.
**D;** The panel shows a volcano plot illustrating the $\log_2$-fold change in transcript abundance versus the adjusted p-value for the Δ*prpC* mutant grown in MOPS-succinate medium compared with the wild-type grown in the same medium. The transcripts that are significantly (q-value < 0.05) increased (red) or decreased (blue) in abundance are indicated. Notable transcripts are labelled (*bkdA1*, *cysI*, *cysD*).
**E;** The panel shows a volcano plot illustrating protein $\log_2$-fold change in transcript abundance versus adjusted p-value for the Δ*prpC* mutant grown in MOPS-succinate + propionate versus the wild-type grown in the same medium. Transcripts which are significantly (q-value < 0.05) increased (red) or decreased (blue) in abundance are indicated. Notable ORFs are labelled (PA3619, *prpB*, *metE*, PA2603, *prpR*, *prpF*, *acnD*, PA3391).
**F;** Western blot showing PrpB (32.1 kDa) protein expression levels in PAO1, the Δ*prpC* mutant and the Δ*acsA* mutant after exposure to increasing concentrations (0 mM, 2 mM, 4 mM) of propionate for 3 h. Isocitrate dehydrogenase (ICD – 45.6 kDa) served as a loading control. Data representative of three independent experiments.
**G-I;** Growth of the wild-type (PAO1), a Δ*prpR* mutant, a Δ*crc* mutant, a Δ*cbrB* mutant, a Δ*acsA* mutant, and a Δ*erdR* mutant in MOPS-succinate (G), MOPS-propionate (H) and MOPS-acetate (I) media. The data are representative of three independent experiments, each performed in triplicate.
**J;** Western blot showing PrpB and ICD expression levels in the indicated mutants (Δ*prpC*, Δ*crc*, Δ*cbrB*, Δ*pycA*, Δ*erdR*) grown in the presence and absence (as indicated) of propionate. Data representative of two independent experiments.

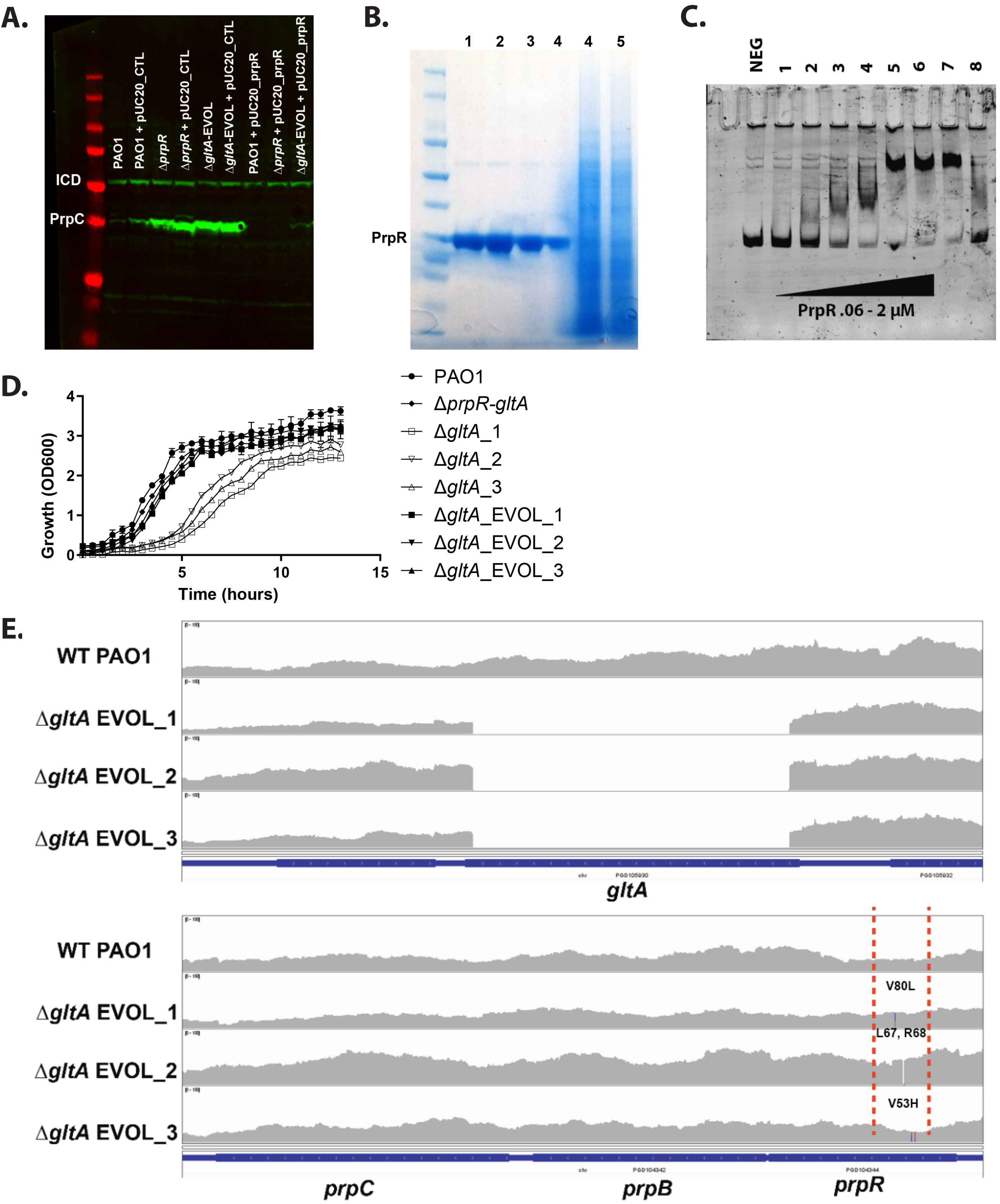


**Figure S6; PrpR functional elucidation and genomic sequencing of the "evolved" Δ*gltA* mutants, Δ*gltA*_EVOL_1-3.**

**A;** Western blot demonstrating that complementation of the Δ*prpR* mutant and the Δ*gltA*_EVOL mutant with a plasmid over-expressing *prpR* (pUCP20_*prpR*) reverts PrpC expression to levels seen in the wild-type in the absence of added propionate. Note that introduction of an empty vector (pUCP20_CTL) into these mutants does not affect PrpC expression compared with the corresponding vector-less mutants. The cultures were all grown in LB medium. Isocitrate dehydrogenase (ICD) served as a loading control. The data are representative of three independent experiments.

**B;** SDS-PAGE analysis showing recombinant expression and purification of His6-PrpR (≈27 kDa) in *E. coli*. Lanes 1-4 = eluted protein fractions from Ni-NTA affinity chromatography. Lane 4 = nickel-column flow through. Lane 5 = whole cell lysate.

**C;** Electrophoretic mobility shift assay (EMSA) polyacrylamide gel showing binding of recombinant PrpR to a PCR-amplified segment of DNA containing the *prpR* promoter labelled with the 6FAM (5 pM), resulting in a protein-DNA complex (lanes 5-7). Lanes 1-6 = increasing PrpR concentration – 0.06 µM, 0.125 µM, 0.25 µM, 0.5 µM, 1 µM, 2 µM. Lane 7 = non-specific competitor (*ccoN1* promoter) in the presence of 2 µM PrpR. Lane 8 = unlabelled competitor (the PCR amplicon containing the *prpR* promoter but not 6FAM-labelled) in the presence of 2 µM PrpR. NEG = negative control (no PrpR added). The data are representative of three independent experiments.

**D;** Growth of PAO1 compared with the Δ*gltA*, Δ*prpR* Δ*gltA* and Δ*gltA*_EVOL mutants in LB medium. The data are representative of three independent experiments, each performed in triplicate.

**E;** Integrative Genomics Viewer (IGV) track highlighting the *gltA* (top) and *prpR-prpC* (bottom) genomic regions in wild-type PAO1 and in the Δ*gltA*_EVOL_1-3 mutants. Reads corresponding to *gltA* are absent in the Δ*gltA*_EVOL_1-3 mutants (as expected), and the single nucleotide polymorphisms and deletions in *prpR* are highlighted (between the red dashed lines).
