## Supplementary material for "Systems-wide dissection of organic acid assimilation in *Pseudomonas aeruginosa* reveals a novel path to underground metabolism": Table_S1

**Table S1A.** Oligonucleotide primers used in this study.

| **Primer Name** | **Sequence (5’ to 3’)** |
| --- | --- |
| *prpC* KO UP F | acccggggatcctctACTCCTTCGTGATCATGGCG |
| *prpC* KO UP R | tgttggcGGCTGTCTGACCGGCCAC |
| *prpC* KO DW F | agacagccGCCAACAACCGCATCATC |
| *prpC* KO DW R | ctgcaggtcgactctGACGATCAGTTGCACCGG |
| *acsA* KO UP F | tcggtacccggggatcctctCTGCTGCTCGGGGTGATC |
| *acsA* KO UP R | cgatcaggtgGTACAGGGATGCCGCAGAC |
| *acsA* KO DW F | atccctgtacCACCTGATCGAGACCCATCG |
| *acsA* KO DW R | catgcctgcaggtcgactctTCGGCCTTGGGATCGTTG |
| *PA3234* KO UP F | acccggggatcctctGGTGATGATCAAGACCGACG |
| *PA3234* KO UP R | gatgaactgGGCAAGGAGACGAGCGATC |
| *PA3234* KO DW F | tccttgccCAGTTCATCCGTTCCCAG |
| *PA3234* KO DW R | ctgcaggtcgactctGGGTAGCGATCAGGGTGTC |
| *PA3568* KO UP F | tcggtacccggggatcctctCGACATCGCATTCCTCGG |
| *PA3568* KO UP R | cctcgatctcGTACCAGGCCAGGGAACG |
| *PA3568* KO DW F | ggcctggtacGAGATCGAGGCTGCGCTG |
| *PA3568* KO DW R | catgcctgcaggtcgactctAAGGTGCTGGGCTGGGAC |
| *aceA* KO UP F | acccggggatcctctCCTACGGGCAGATCTACAG |
| *aceA* KO UP R | tctttgccAACTGCCTTGATCTCGTTC |
| *aceA* KO DW F | aggcagttAACCAGTTCCACTAAGGAACAG |
| *aceA* KO DW R | ctgcaggtcgactctGTCGCCGGAGTAGCAGAC |
| *glcB* KO UP F | acccggggatcctctTGTAGCCCTTCCACCAGTTC |
| *glcB* KO UP R | tcttcgcCAGGCCACCGACTTGAACs |
| *glcB* KO DW F | gtggcctgGCGAAGAACGGGCTGTAG |
| *glcB* KO DW R | ctgcaggtcgactctGCAGGCCGAGGACATGAG |
| *mmsB* KO UP F | acccggggatcctctTTCATCCTCAAGCCTTCCGAG |
| *mmsB* KO UP R | cttctggatCAGGCCGAGGAATGCGATG |
| *mmsB* KO DW F | tcggcctgATCCAGAAGCTCTACCGCG |
| *mmsB* KO DW R | ctgcaggtcgactctCACATCAGCGGGCCGTAG |
| *prpR* KO UP F | acccggggatcctctGGCACGGTGGAAGAGAAG |
| *prpR* KO UP R | gaagcggaGTCATGGGCAATCTCGAG |
| *prpR* KO DW F | cccatgacTCCGCTTCCTCCACCGTC |
| *prpR* KO DW R | ctgcaggtcgactctTTTCTGGATGCTGCAACTCG |
| *cbrB* KO UP F | acccggggatcctctAAGACCTCACCGAAACCC |
| *cbrB* KO UP R | cgcggaatGACGATCAGAATATGTGCCATG |
| *cbrB* KO DW F | gatcgtcATTCCGCGGCGTAAATC |
| *cbrB* KO DW R | ctgcaggtcgactctGCGGCATTATCAGGACC |
| *crc* KO UP F | acccggggatcctctACGCATTACATCCTGTCC |
| *crc* KO UP R | gggctcaGTTCACACTGATGATCCGC |
| *crc* KO DW F | gtgtgaacTGAGCCCTGTCAGACACG |
| *crc* KO DW R | ctgcaggtcgactctCCGCACCGCCTCAATCAC |
| *gltA* KO UP F | tcggtacccggggatcctctGGTCAACAGGCTGAAGATG |
| *gltA* KO UP R | cggtgaagtcTGAGCCCTCGATGATCAAC |
| *gltA* KO DW F | cgagggctcaGACTTCACCGCCCTCAAG |
| *gltA* KO DW R | catgcctgcaggtcgactctGAAAATCATCTCTGCGGG |
| *pEX19Gm* F | AGAGTCGACCTGCAGGCATG |
| *pEX19Gm* R | AGAGGATCCCCGGGTACC |
| *acsA* *Tn7T* *lux* F *BamHI* | AAACGCGGATCCCCTGGGCATCGTCATGGT |
| *acsA* Tn7T lux R *XhoI* | AAAACTCGAGCACGGGGTACAGGGATGC |
| *prpC pet19m F NdeI* | AAAAAACATATGATGGCTGAAGCAAAAGTACTGAG |
| *prpC pet19m R XhoI* | CGCGGATCCTCAGCGTTGCTCCAGCG |
| *gltA pet19m F NdeI* | AAAAAACATATGGCTGACAAAAAAGCGCAGTT |
| *gltA pet19m R XhoI* | ACGCGGATCCTCAGCCGCGATCCTTGAG |
| *prpR_pUCP20_F* | tcaaggagtcgaaagcATGATGACCCAGAACCCGCCC |
| *prpR_pUCP20_R* | cgacggccagtgccaTTAGACGGTGGAGGAAGCG |
| *prpRp_F* | CTGCTTCATCACGTCGCC |
| *prpRp_R* | GTCATGGGCAATCTCGAGAG |

**Table S1B:** Bacterial strains and plasmids used in this study

| **Strain or plasmid** | **Source** |
| --- | --- |
| **Strains** |  |
| *E. coli* JM109 | New England Biolabs |
| *P. aeruginosa* PAO1 | (1) |
| PAO1 *ΔprpC* | This study |
| PAO1 *ΔacsA* | This study |
| PAO1 *ΔaceA* | This study |
| PAO1 *ΔglcB* | This study |
| PAO1 *ΔPA3234* | This study |
| PAO1 *ΔmmsB* | This study |
| PAO1 *ΔPA3568* | This study |
| PAO1 *ΔgltA* | This study |
| PAO1 *aceA::lux* | (2) |
| PAO1 *glcB::lux* | (2) |
| PAO1 *cco2::lux* | (2) |
| pycA (PA5435)::PW10177 | (3) |
| pycB ( PA5436)::PW10180 | (3) |
| **Plasmids**  pEX19Gm (*P. aeruginosa* suicide vector, Gm) | (4) |
| pUC18T-mini-Tn7T-lux-Gm (mini-Tn7 luxCDABE transcriptional fusion vector) | (5) |
| Pet19m | (6) |
| pUCP20 | (7) |

**Table S1C:** The steady-state kinetic parameters of PaPrpC and several other PrpC from different organisms.

| Protein | Organism | *K_m_* (OAA, µM) | *k_cat/_K_m_* (OAA) | *K_m_* (PrCoA, µM) | *k_cat/_K_m_* (PrCoA) | *K_m_* (AcCoA, µM) | *k_cat/_K_m_* (AcCoA) | Reference |
| --- | --- | --- | --- | --- | --- | --- | --- | --- |
| PrpC | *P. aeruginosa* | 15.65**^a^** | 228 x 10^3^ M^-1^s^-1^ | 45 | 104 x 10^3^ M^-1^s^-1^ | 28 | 114 x 10^3^ M^-1^s^-1^ | This work |
| PrpC | *P. aeruginosa* | 2**^a^** | N/A | N/A | N/A | 50 | N/A | (8) |
| PrpC | *S. enterica* | 15^a^ & 13^b^ | N/A | 45 | 183 x 10^-3^ M^-1^s^-1^ | 265 | 7 x 10^-3^ M^-1^s^-1^ | (9) |
| PrpC | *S. enterica* | 14^a^ & 12^b^ | N/A | 48 | 150 x 10^3^ M^-1^s^-1^ | 285 | 5 x 10^3^ M^-1^s^-1^ | (10) |
| PrpC | *E. coli*^c^ | 5* | N/A | 37 | N/A | 101 | N/A | (11) |
| PrpC | *E. coli* | N/A | N/A | 17 | N/A | N/A | N/A | (12) |
| MmgD^d^ | *B. Subtilis* | N/A | N/A | N/A | 41 x 10^3^ M^-1^s^-1^ | N/A | 18 x 10^3^ M^-1^s^-1^ | (13) |
| CS | DS2-3R | 3^a^ & 7^b^ | N/A | 16 | 52 x 10^4^ M^-1^s^-1^ | 229 | 9 x 10^4^ M^-1^s^-1^ | (11) |

**^a^***K_m_* determined by fixed concentration of PrCoA (200 µM)

**^b^***K_m_* determined by fixed concentration of AcCoA (200 µM)

**^c^**The activity was measured using cell extracts.

^d^MmgD is the equivalent of PrpC in *B. Subtilis*

**K_m_* value for both acyl-CoAs

**Table S****1D.** Crystallographic statistics for PrpC-Apo, PrpC-oxaloacetate bound, and GltA-Apo.

| Structure | PrpC-Apo | PrpC-Oxaloacetate bound | GltA-Apo |
| --- | --- | --- | --- |
| **PDB ID Code** | **6S6F** | **6S87** | **6ZU0** |
| **Data Collection** |  |  |  |
| Wavelength (Å) | 0.9795 | 0.9282 | 0.9762 |
| Resolution range (Å) | 49.91 - 1.53 (1.585 - 1.53) | 25.78 - 1.65 (1.709 - 1.65) | 73.5 - 3.397 (3.519 - 3.397) |
| Space group | P 1 2_1_ 1 | P 1 2_1_ 1 | C 1 2 1 |
| Unit cell |  |  |  |
| *a, b, c* (Å) | 51.01 83.19 82.71 | 51.64 86.76 161.84 | 216.74 80.17 196.65 |
| *a, b, g* (°) | 90 101.92 90 | 90 95.89 90 | 90 121.88 90 |
| Total reflections | 329742 (23523) | 708405 (53561) | 173880 (8498) |
| Unique reflections | 101634 (10050) | 170219 (16998) | 39614 (3854) |
| Multiplicity | 3.2 (3.1) | 4.2 (4.3) | 4.5 (4.4) |
| Completeness (%) | 98.92 (97.07) | 99.67 (99.78) | 99.01 (97.40) |
| Mean I/sigma(I) | 8.9 (1.2) | 11.5 (1.1) | 4.7 (1.3) |
| Wilson B-factor | 18.82 | 24.05 | 87.09 |
| R-merge | 0.056 (0.936) | 0.064 (1.362) | 0.176 (0.895) |
| R-meas | 0.067 (1.128) | 0.073 (1.557) | 0.225 (1.153) |
| R-pim | 0.036 (0.622) | 0.035 (0.747) | 0.139 (0.715) |
| CC1/2 | 0.998 (0.562) | 0.999 (0.578) | 0.988 (0.566) |
| **Refinement** |  |  |  |
| Resolution range (high resolution) (Å) | 49.91 - 1.53 (1.585 - 1.53) | 25.78 - 1.65 (1.709 - 1.65) | 73.5 - 3.39 (3.51 - 3.39) |
| Reflections used in refinement | 100694 (9823) | 169968 (16972) | 39586 (3854) |
| Reflections used for R-free | 5036 (493) | 8364 (843) | 1894 (201) |
| R-work | 0.1971 (0.3519) | 0.2130 (0.3578) | 0.2383 (0.3640) |
| R-free | 0.1965 (0.3565) | 0.2130 (0.3608) | 0.2654 (0.4346) |
| Number of non-hydrogen atoms | 6121 | 11690 | 19923 |
| Macromolecules | 5793 | 11287 | 19923 |
| Ligand | 37 | 51 | - |
| Solvent | 291 | 352 | - |
| Protein residues | 730 | 1431 | 2543 |
| RMS (bonds) Å | 0.016 | 0.017 | 0.007 |
| RMS (angles) | 1.48 | 1.8 | 1.31 |
| Ramachandran favoured (%) | 98.35 | 98.38 | 95.09 |
| Ramachandran allowed (%) | 1.65 | 1.55 | 4.35 |
| Ramachandran outliers (%) | 0 | 0.07 | 0.55 |
| Rotamer outliers (%) | 0.82 | 1.27 | 3.19 |
| Clashscore | 2.41 | 1.33 | 8.15 |
| Average B-factor | 26.35 | 31.88 | 92.93 |
| Macromolecules | 26.04 | 31.84 | 92.93 |
| Ligands | 39.64 | 36.87 | - |
| Solvent | 30.77 | 32.55 | - |

1.	Stover CK, Pham XQ, Erwin AL, Mizoguchi SD, Warrener P, Hickey MJ, Brinkman FSL, Hufnagle WO, Kowalik DJ, Lagrou M, Garber RL, Goltry L, Tolentino E, Westbrock-Wadman S, Yuan Y, Brody LL, Coulter SN, Folger KR, Kas A, Larbig K, Lim R, Smith K, Spencer D, Wong GK-S, Wu Z, Paulsen IT, Reizer J, Saier MH, Hancock REW, Lory S, Olson M V. 2000. Complete genome sequence of Pseudomonas aeruginosa PAO1, an opportunistic pathogen. Nature 406:959–964.

2.	Dolan SK, Kohlstedt M, Trigg S, Vallejo Ramirez P, Kaminski CF, Wittmann C, Welch M. 2020. Contextual Flexibility in *Pseudomonas aeruginosa* Central Carbon Metabolism during Growth in Single Carbon Sources. MBio 11.
